# Integrated gene analyses of *de novo* mutations from 46,612 trios with autism and developmental disorders

**DOI:** 10.1101/2021.09.15.460398

**Authors:** Tianyun Wang, Chang Kim, Trygve E. Bakken, Madelyn A. Gillentine, Barbara Henning, Yafei Mao, Christian Gilissen, The SPARK Consortium, Tomasz J. Nowakowski, Evan E. Eichler

## Abstract

Most genetic studies consider autism spectrum disorder (ASD) and developmental disorder (DD) separately despite overwhelming comorbidity and shared genetic etiology. Here we analyzed *de novo* mutations (DNMs) from 15,560 ASD (6,557 are new) and 31,052 DD trios independently and combined as broader neurodevelopmental disorders (NDD) using three models. We identify 615 candidate genes (FDR 5%, 189 potentially novel) by one or more models, including 138 reaching exome-wide significance (p < 3.64e-07) in all models. We find no evidence for ASD-specific genes in contrast to 18 genes significantly enriched for DD. There are 53 genes show particular mutational-bias including enrichments for missense (n=41) or truncating DNM (n=12). We find 22 genes with evidence of sex-bias including five X chromosome genes also with significant female burden (*DDX3X, MECP2, SMC1A, WDR45*, and *HDAC8)*. NDD risk genes group into five functional networks associating with different brain developmental lineages based on single-cell nuclei transcriptomic data, which provides important insights into disease subtypes and future functional studies.

## INTRODUCTION

Neurodevelopmental disorders (NDDs) are a group of heritable disorders that are twice as likely to affect males when compared to females and whose prevalence broadly defined continues to increase, in part, due to improved pediatric ascertainment^1^. Among NDDs, autism spectrum disorder (ASD), developmental disorder (DD), intellectual disability (ID), and attention deficit hyperactivity disorder (ADHD) show considerable heterogeneity both genetically and clinically, and often present some of the greatest sex differences in prevalence^2-6^. Individuals with different primary NDD diagnoses frequently present overlapping phenotypes. For example, ADHD has been observed as the most common co-occurring disorder among individuals with an ID diagnosis, followed by 15-40% of individuals presenting with ASD^7-10^. Similarly, ∼30% of individuals with an ASD diagnosis also show some level of cognitive impairment, and ∼30-40% of cases co-occur with ADHD^5,11^. This significant diagnostic overlap among NDD groups has long suggested a shared genetic etiology, at least among a subset of cases^12^.

*De novo* mutations (DNMs) that disrupt or alter protein-coding gene function significantly contribute to NDDs, this genetic signal has facilitated the discovery of hundreds of risk genes over the last decade^13,14^. Most studies to date have focused on a single phenotype group, e.g., either on ASD^15^ or DD^16^, in an effort to identify genes that specifically contribute to that phenotype. The locus heterogeneity of these disorders, however, has limited power to identify genes with rare DNMs given that more than half of the genes still await a sufficient number of cases to reach statistical significance^17^. Moreover, different studies have tended to apply their preferred statistical or evolutionary models to identify genes with an excess of DNM. This has led to the emergence of slightly different gene sets from the same parent–child trio data^15,17^.

In this study, we integrate DNMs from 11 cohorts with a primary diagnosis of ASD or DD from 46,612 parent–child trios, of which 6,557 ASD trios are new to this study, and apply three different statistical models (chimpanzee–human divergence [CH] model^17,18^, denovolyzeR^19^, and DeNovoWEST^16^) (Figure 1). Briefly, the CH model estimates the number of expected DNMs by incorporating locus-specific transition, transversion, and indel rates, the gene length, and null expectation based on chimpanzee–human coding sequence divergence; denovolyzeR estimates mutation rates, considers the triplet context, and adjusts divergence based on macaque–human gene comparisons; DeNovoWEST applies the same underlying mutation rate as denovolyzeR but also incorporates gene-based weighting with missense clustering to provide the most sensitive set of candidate genes. In addition to applying standard statistical thresholds of significance, we prioritized genes as higher confidence if they were predicted by more than one or all of these three models and flag potential artefacts in previous call sets based on exclusivity of variant calls to a particular study.

**Figure 1.**
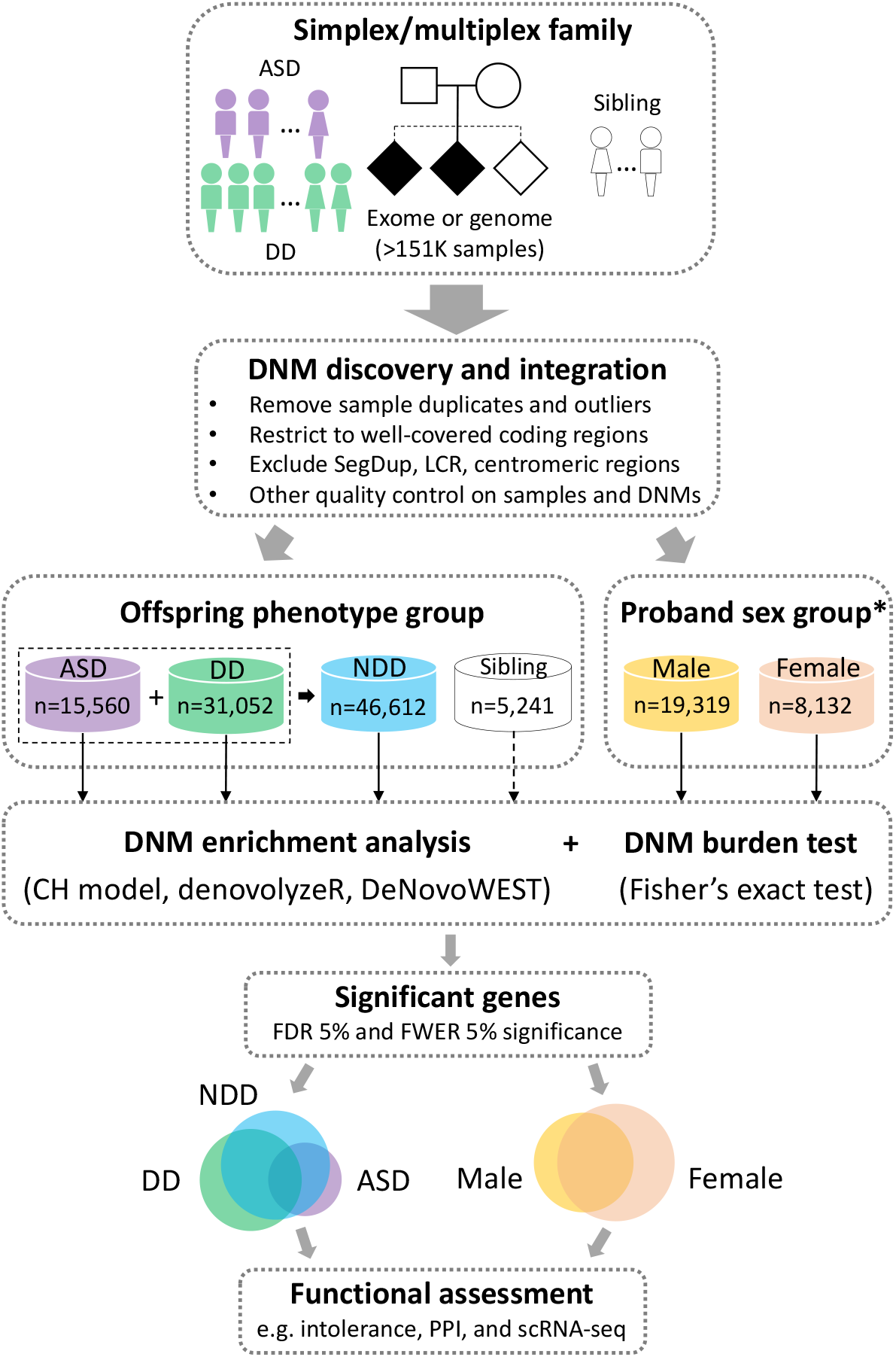
Study workflow. DNMs from >151K samples including both simplex and multiplex families with a primary diagnosis of ASD or DD were integrated with strict quality control and filtering measures applied (Online Methods). *De novo* enrichment analysis was performed independently in ASD (n = 15,560), DD (n = 31,052), and NDD (n = 46,612) groups, and also a subset of probands with sex information available were grouped by males and females, in parallel using three statistical models (CH model, denovolyzeR, and DeNovoWEST). Siblings were also analyzed using the CH model and denovolyzeR, but not run for DeNovoWEST due to the small sample size (n = 5,241). Significant genes were used for downstream analyses for the identification of new risk genes and the comparison between phenotype and sex. *Sex information is only available for a subset (58.9%, 27,451/46,612) of the probands.

The goal of this analysis is fourfold: 1) discover new candidate genes by combining ASD and DD patients as a broader NDD cohort; 2) identify ASD- or DD-enriched genes and test if any genes show an enrichment in one diagnosis over the other^20^; 3) identify genes that favor a particular class of mutation (i.e., missense over likely gene-disruptive [LGD]); and 4) identify genes that are biased for DNMs either among females or males. We further explore the protein-protein interaction (PPI) networks and expression properties of these genes both in bulk tissue as well as single-cell RNA-seq data to reveal novel properties regarding the tissues and functional neural networks affected by disruption of these genes.

## RESULTS

### DNM integration across studies

We integrated whole-exome sequencing (WES) and whole-genome sequencing (WGS) data from over 44,800 parent–child families from 11 cohorts^15,16,21-26^, where children received a primary diagnosis of ASD or DD (Table S1). We first split the cohorts into two groups based on whether the underlying Illumina sequencing data were available for reanalysis and variant recalling. We identified DNMs by reanalyzing underlying CRAMs or BAMs using a previously published pipeline^27^ using FreeBayes and GATK for five of the cohorts (recalled subset: 46.4% of families or 60,868 exomes and 9,304 genomes). For the remaining six cohorts (no-recall subset: 53.6% of families with 81,052 exomes), we used the published DNMs with downstream filtering applied (Table S2). We performed stringent quality control and filtering on both recalled and no-recall subsets in an effort to better harmonize DNMs (Methods). This included removing sample duplicates and outliers with a significant excess of DNM; restricting calls to regions with sufficient coverage (∼20X) across the different platforms, including WES and WGS data; excluding DNMs within segmental duplications, low complexity regions, and centromeric and telomeric regions to reduce false positives; and flagging “jackpot” genes where multiple independent events were identified in a single cohort but never again observed in another study (Methods).

After quality control on both samples and variants, the harmonized DNM set was generated from 15,560 ASD (6,557 are new) and 31,052 DD patients as well as 5,241 unaffected siblings (3,034 are new). This set includes 6,921 *de novo* LGD variants (dnLGD, including frameshift, stop-gain, splice-donor, or splice-acceptor variant), 32,774 *de novo* missense variants (dnMIS), as well as 11,706 *de novo* synonymous variants (dnSYN). Among the dnMIS variants, we classify 5,946 as severe based on a combined annotation dependent depletion (CADD) score^28,29^ greater than 30 (v1.3), ranking these mutations among the top 0.1% for the most severe predicted effect (dnMIS30) (Table S3). There are 11,409 genes with non-synonymous DNMs in probands versus 2,742 genes in siblings, with 37.7% (4,304/11,409) of the genes in probands and 83.7% (2,296/2,742) of genes in siblings with only one DNM (Figure S1). Comparing the overall mutation rate for the recalled and no-recall sample sets gave similar DNM frequency (∼0.78 non-synonymous DNMs per proband). Considering cohorts by phenotype, we find that DD shows the highest DNM rate followed by ASD and then siblings (Figure S2, Table S2).

### DNM-enriched NDD candidate genes and diagnostic specificity

To identify genes with a significant excess of DNMs, we applied three statistical models (CH model^17,18^, denovolyzeR^19^, and DeNovoWEST^16^) independently to the ASD and DD cohorts based on their primary diagnoses and then combined as one broader NDD group (Figure 1). After excluding genes that showed evidence of DNM significance among unaffected siblings (n = 7, Table S4), we identify 615 candidate genes in the combined NDD group with an excess of DNMs by one or more of the three models (union FDR 5%, q-value < 0.05, DNM count > 2). We consider these candidates as a lower confidence set and hereafter refer as the LC615 gene set (Table 1, Figure 2). Although the majority of the LC615 genes are shared across the models, 43.3% (266/615) are only identified by a single model (Figure S3). Among the LC615 genes, we also defined the highest confidence set of 138 genes (HC138 genes) where the excess of DNMs meets exome-wide significant supported by all three models (intersection family-wise error rate [FWER] 5%, p-value < 3.64e-07, DNM count > 2) (Table 1, Table S5). Among the three models, DeNovoWEST shows the greatest sensitivity because it incorporates missense clustering in addition to strict counts of DNMs^16^. We observe similar trends in our independent analyses of the ASD and DD cohorts (Table 1, Tables S6-S7). Candidate genes, as expected, are significantly more likely to be intolerant to mutation (Figure S4). Among the LC615 genes, 324 genes are considered novel when compared to three previous reports (ASC102^15^, Coe253^17^, DDD285^16^) (Figure S5, Table S8). Among the 324 candidates with novel statistical significance, we searched for additional evidence of pathogenicity (including case reports) within the Development Disorder Genotype - Phenotype Database (DDG2P), SFARI gene, Online Mendelian Inheritance in Man (OMIM) database, and PubMed, identifying 189 genes (30 FWER genes) with novel association with NDD (Table S8). Among the most stringent set of HC138 genes, we identify or confirm three genes (*MED13, NALCN*, and *PABPC1*) newly reach exome-wide significance in this large data, further confirmed their association with NDD based on recent reports^30,31^ (Figure S5). We should also note that there are 52 potentially novel genes if we consider exome-wide significance more broadly by any one or more of the models instead of requiring all three to intersect (Table S8). These genes, thus, represent a powerful resource for future clinical and basic research investigations related to NDDs.

**Table 1.** Genes with significant excess of DNM across phenotype and sex groups.

| Group | Samples | # DNMs |  |  | FDR 5% significant genes (q-value < 0.05) |  |  |  |  | FWER 5% significant genes (p-value < 3.64e-07) |  |  |  |  |
| --- | --- | --- | --- | --- | --- | --- | --- | --- | --- | --- | --- | --- | --- | --- |
|  |  | dnLGD | dnMIS | dnMIS30 | CH model | denovolyzeR | DeNovoWest | Union | Intersection | CH model | denovolyzeR | DeNovoWest | Union | Intersection |
| Offspring phenotype group |  |  |  |  |  |  |  |  |  |  |  |  |  |  |
| ASD | 15,560 | 1,661 | 9,187 | 1,454 | 70 | 52 | 113 | 133 | 41 | 26 | 22 | 27 | 39 | 16 |
| DD | 31,052 | 4,931 | 20,588 | 4,084 | 352 | 293 | 414 | 511 | 233 | 181 | 166 | 196 | 241 | 136 |
| NDD* | 46,612 | 6,592 | 29,775 | 5,538 | 399 | 323 | 479 | 615 (LC615) | 237 | 208 | 171 | 190 | 264 | 138 (HC138) |
| Sibling | 5,241 | 329 | 2,999 | 408 | 3 | 6 | - | 7 | - | 1 | 1 | - | 2 | - |
| Proband sex group |  |  |  |  |  |  |  |  |  |  |  |  |  |  |
| Male | 19,319 | 2,262 | 11,714 | 1,990 | 131 | 106 | 168 | 207 | 78 | 60 | 54 | 58 | 83 | 35 |
| Female | 8,132 | 1,371 | 5,287 | 1,014 | 124 | 114 | 159 | 177 | 94 | 63 | 52 | 70 | 83 | 42 |

Two levels of significance, FDR 5% and FWER 5%, were assessed based on the union and intersection of three statistical models. The lower confidence set (LC615) is based on the union of all genes at FDR 5% significance observed in any of the three models; the highest confidence set (HC138) is based on the intersection of genes with FWER 5% significance supported by all three models. The FDR 5% significance threshold was corrected by the Benjamini–Hochberg method for all genes in each model (18,946 genes in CH model, 19,618 genes in denovolyzeR, and 18,762 genes in DeNovoWEST); the FWER 5% significance threshold was corrected by Bonferroni method for 19,618 genes (the largest among three models) and seven tests in the analyses (tests for dnLGD, dnMIS, and dnMIS30 variants in CH model; dnLGD and dnMIS variants in denovolyzeR; and DNM and dnMIS variants in DeNovoWEST). Genes with DNM significance in siblings were excluded from all sets (n=7). \*NDD = ASD + DD. Details regarding the NDD risk genes can be found in Tables S4-S7, S12, S13.

**Figure 2.**
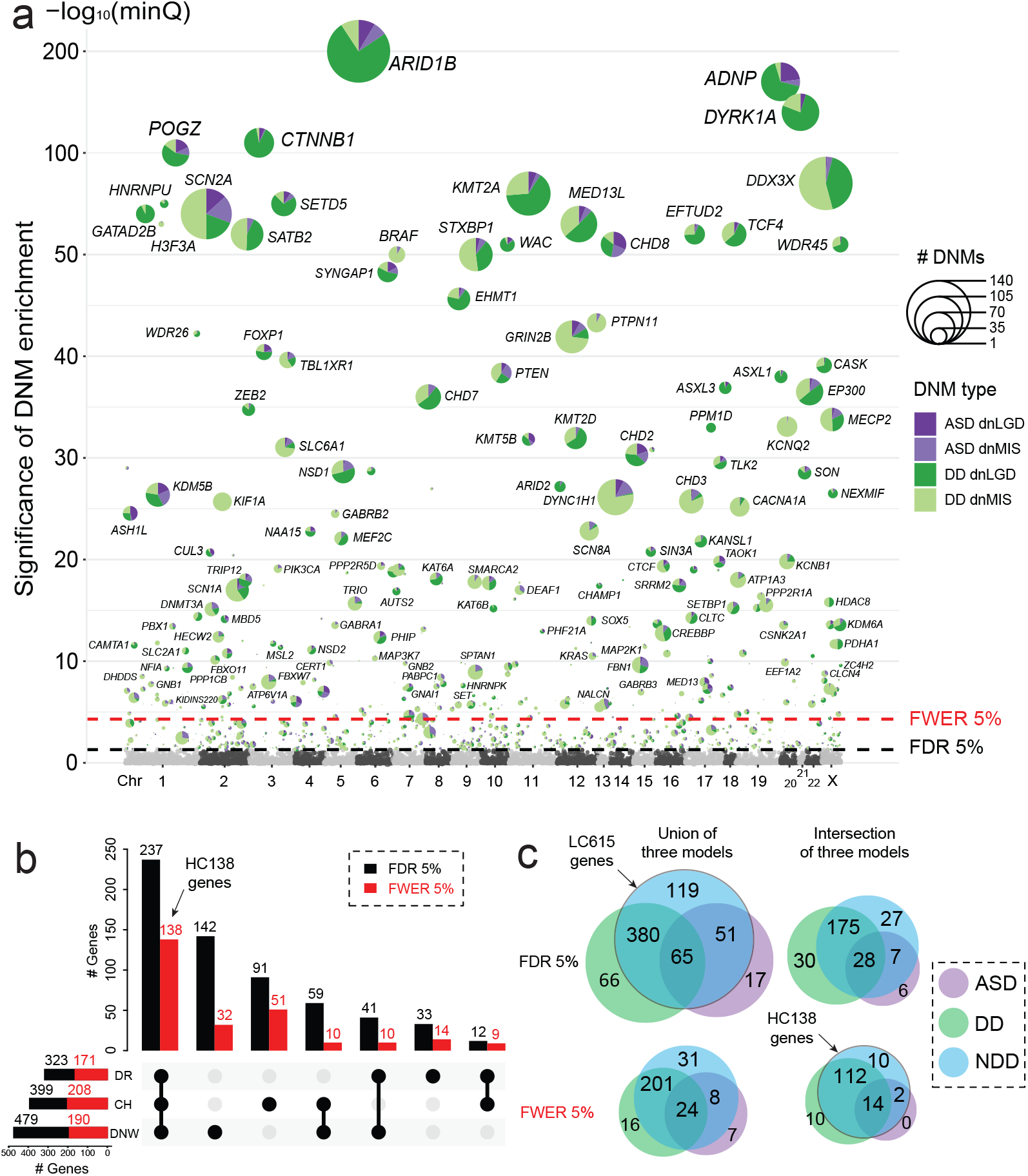
*De novo* enrichment analysis and significant genes by model and phenotype. **(a)** The smallest q-value (minQ) after Benjamini–Hochberg correction of each gene across the three models was plotted in alphabetic order of gene name by chromosome. The LC615 genes reaching union FDR 5% significance were plotted with the number of dnLGD and dnMIS variants in ASD and DD scaled in pie chart, the HC138 genes reaching the intersection FWER 5% significance were additionally labeled with gene name. **(b)** UpsetR plot shows the number of genes that reach FDR 5% (black bar) and FWER 5% (red bar) significance identified by each of the three models (DR: denovolyzeR; CH: CH model; DNW: DeNovoWEST) in the combined NDD set. **(c)** The overlap FDR 5% and FWER 5% significant genes across phenotype groups (ASD, DD, and NDD) at both the union and intersection of the three models. The union FDR 5% set are the LC615 genes, and the intersection FWER 5% set are the HC138 genes.

Next, we considered cohorts with the primary diagnosis of ASD and DD separately using the same criteria and then compared the groups to identify genes potentially unique to a specific diagnosis. This analysis identifies several genes enriched for DNMs that are unique to each primary diagnosis or combined (union FDR 5%): ASD (n = 17), DD (n = 66), and NDD (n = 119) (Figure 2c). However, most of those genes are only identified by a single model and often with borderline statistical support (88.2% [15/17] of genes in ASD, 93.9% [62/66] of genes in DD, and 85.7% [102/119] of genes in the NDD group). Among the genes with unique significance in ASD, we note that 58.8% (10/17) of genes show evidence of DNMs in both ASD and DD patients, which suggesting that screening of further DD samples will likely yield additional cases and eliminate the ASD specificity. Although there are seven genes (*INTS2, ASB17, KRT34, C1orf105, C6orf52, NUCB2*, and *JKAMP*) that currently show no DNMs among the DD patients, but none are supported by all three models (five of which only predicted by DeNovoWEST) and none reach exome-wide significance in ASD, and most are not particularly compelling candidates based on available functional data. We suggest that all of these should be regarded as either unlikely or low-confidence “autism-specific” genes. While we recognize that there is considerable overlap among DD and autism, autism criteria are generally more specific including a large number of patients without intellectual disability or generalized developmental delay.

In contrast to ASD, there is compelling evidence for DD-specific genes that are not associated or rarely associated with ASD. Among the 66 genes that reach FDR significance only in DD but neither ASD nor the combined NDD set, 10 genes meet the exome-wide significance and supported by all three models (*ARID1A, BCOR, PTCH1, KIF11, DPF2, DNM1L, ANKRD11, H4C5, HUWE1*, and *TRAF7*) (Figure 2c). There are three genes (*KIF11, H4C5*, and *TRAF7*) that show DD-specific significance where no DNM has yet been observed among autism patients in this study, however, there are also few autism cases have been reported previously with DNM in *KIF11*^*32*^, and *TRAF7* ^*33*^. As expected, genes with NDD-specific significance show DNMs in both ASD and DD patients, but 10 genes reach the intersection exome-wide significance when cohorts are combined together as broader NDD group (*MYT1L, PHF21A, FBN1, PBX1, GNB2, PABPC1, CLCN4, NALCN, PSMC5*, and *CERT1*).

Comparing the relative frequency of DNMs among DD and ASD cohorts identifies genes with a potential bias toward DD or ASD diagnoses, especially when DNM is further categorized by dnMIS or dnLGD mutation class (Figure 3, Tables S9-S10). There are chromatin modifying genes among the genes with a trending of potential ASD bias: *CHD8, KDM5B, ASH1L*, and *KMT5B*—although none of these genes reach significance for ASD enrichment. In contrast to DD patients, a comparison of DNM counts identifies 18 genes with higher DNM burden in DD over ASD patients (two-sided Fisher’s exact test, FDR 5%) (Table 2). Of which, *GATAD2B* and *KIF1A* are exclusive to DD without DNM in ASD patients in this study (Figure 4). Interestingly, *GATAD2B* shows an enrichment only for dnLGD variants, while in contrast *KIF1A* shows a specific enrichment for dnMIS variants (Figure 4). Patients with mutations in these genes have been described as exhibiting DD and moderate to severe ID^34,35^. Although no gene shows significant burden specifically for ASD compared to DD patients, however, there are 149 of the LC615 genes show a trend toward ASD diagnosis over DD with more DNMs in ASD patients when comparing DNM frequency with sample size adjusted. For example, *CHD8* is highly intolerant to LGD mutation (pLI score = 1, LOEUF score = 0.082), shows a 2.22-fold enrichment of DNMs in ASD patients (n = 15,560, 18 dnLGD and 12 dnMIS variants) when compared to DD patients (n = 31,052, 19 dnLGD and 8 dnMIS variants). Other genes, like *KDM5B* (1.48-fold) and *WDFY3* (2.90-fold), also have more DNMs in ASD than in DD patients relative to the corresponding sample size (Figure 4, Table S10).

**Figure 3.**
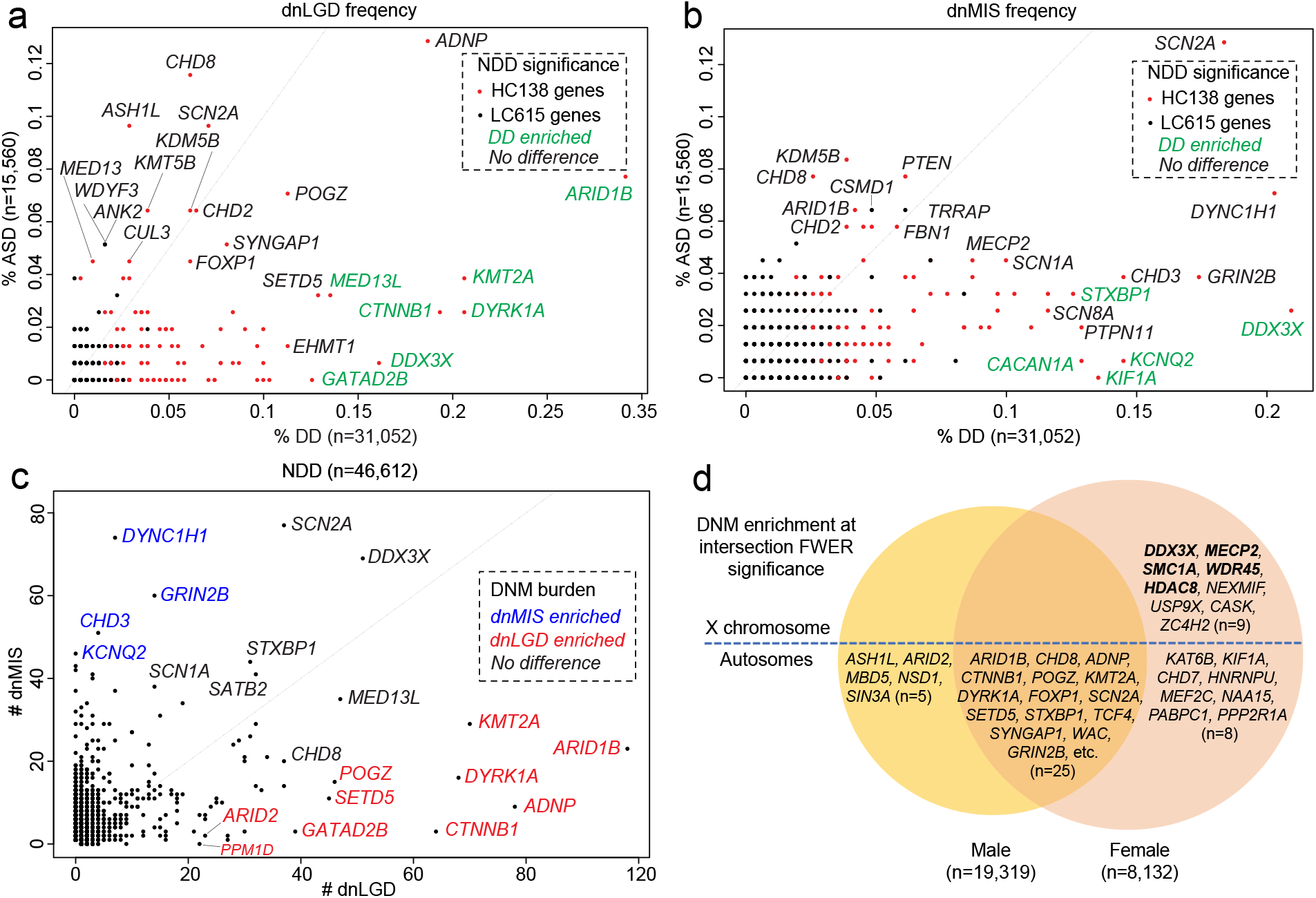
Genes with phenotype, mutation class, and sex-bias DNM burden. **(a)** dnLGD and **(b)** dnMIS frequencies in ASD (y axis) and DD (x axis) patients were plotted for all LC615 genes in the combined NDD group, with the HC138 genes in red and others in black dots. Genes with enriched DNMs in DD over ASD patients were in green. **(c)** Number of dnLGD and dnMIS variants for all genes with DNM in the combined NDD group. Example genes with significant burden of dnLGD (red) or dnMIS (blue) variants compared to the other mutation class are labeled with gene name in color. **(d)** Genes with potential sex bias. Males and females were treated as two separate groups and genes that reached intersection FWER significance (p < 3.64e-07, by all three models) for DNM in males (left) and females (right) as opposed to both sexes (center of Venn). The five genes (bold) with a significant excess of DNM in females (FDR 5%, two-sided Fisher’s exact test) all map to the X chromosome.

**Table 2.** Genes with excess DNM burden in DD versus ASD patients.

| Gene | DD DNM<br>(dnLGD/dnMIS) | ASD DNM<br>(dnLGD/dnMIS) | p-value | q-value | OR [95% CI] | Significant variant |
| --- | --- | --- | --- | --- | --- | --- |
| *ARID1B* | 119 (106/13) | 22 (12/10) | 2.51e-06 | 5.02e-03 | 2.7 [1.7 – 4.5] | DNM&dnLGD |
| *DDX3X* | 115 (50/65) | 5 (1/4) | 5.41e-15 | 1.08e-10 | 11.6 [4.8 – 36.3] | DNM&dnLGD&dnMIS |
| *KMT2A* | 90 (64/26) | 9 (6/3) | 2.25e-08 | 1.50e-04 | 5 [2.5 – 11.3] | DNM&dnLGD |
| *DYRK1A* | 80 (64/16) | 4 (4/0) | 3.11e-10 | 3.11e-06 | 10 [3.8 – 37.8] | DNM&dnLGD |
| *MED13L* | 72 (42/30) | 10 (5/5) | 1.71e-05 | 2.14e-02 | 3.6 [1.9 – 7.9] | DNM |
| *SATB2* | 68 (32/36) | 5 (0/5) | 1.31e-07 | 4.37e-04 | 6.8 [2.8 – 21.7] | DNM&dnLGD |
| *STXBP1* | 67 (28/39) | 8 (3/5) | 7.10e-06 | 1.18e-02 | 4.2 [1 – 10.1] | DNM |
| *CTNNB1* | 62 (60/2) | 5 (4/1) | 7.84e-07 | 1.74e-03 | 6.2 [2.5 – 19.8] | DNM&dnLGD |
| *TCF4* | 51 (31/20) | 4 (3/1) | 9.21e-06 | 1.42e-02 | 6.4 [2.4 – 24.4] | DNM |
| *KMT2D* | 47 (30/17) | 3 (0/3) | 6.63e-06 | 1.18e-02 | 7.9 [2.5 – 39.5] | DNM&dnLGD |
| *KCNQ2* | 45 (0/45) | 1 (0/1) | 3.01e-07 | 8.61e-04 | 22.6 [3.9 – 907.9] | DNM&dnMIS |
| *CACNA1A* | 43 (3/40) | 1 (0/1) | 5.05e-07 | 1.26e-03 | 21.6 [3.7 – 868.6] | DNM&dnMIS |
| *EFTUD2* | 43 (31/12) | 3 (1/2) | 2.67e-05 | 3.14e-02 | 7.2 [2.3 – 36.2] | DNM |
| *WDR45* | 35 (24/11) | 1 (1/0) | 1.06e-05 | 1.51e-02 | 17.6 [3 – 710.9] | DNM |
| *CASK* | 34 (24/10) | 1 (1/0) | 1.71e-05 | 2.14e-02 | 17.1 [2.9 – 691.2] | DNM |
| *GATAD2B* | 42 (39/3) | 0 (0/0) | 5.52e-08 | 2.21e-04 | Inf [5.5 – Inf] | DNM&dnLGD |
| *KIF1A* | 42 (0/42) | 0 (0/0) | 5.52e-08 | 2.21e-04 | Inf [5.5 – Inf] | DNM&dnMIS |
| *CHD7* | 51 (31/20) | 6 (0/6) | 7.36e-06 | 1.64e-02 | Inf [4 – Inf] | dnLGD only |

Two-sided Fisher's exact test for DNM counts was performed comparing ASD (n = 15,560) and DD (n = 31,052) patients. This table shows the 18 genes with significant DNM burden in DD compared to ASD patients, while no gene with significant burden is identified in ASD patients. The significance threshold (FDR 5%) was corrected for 20,000 genes (Benjamini–Hochberg).

**Figure 4.**
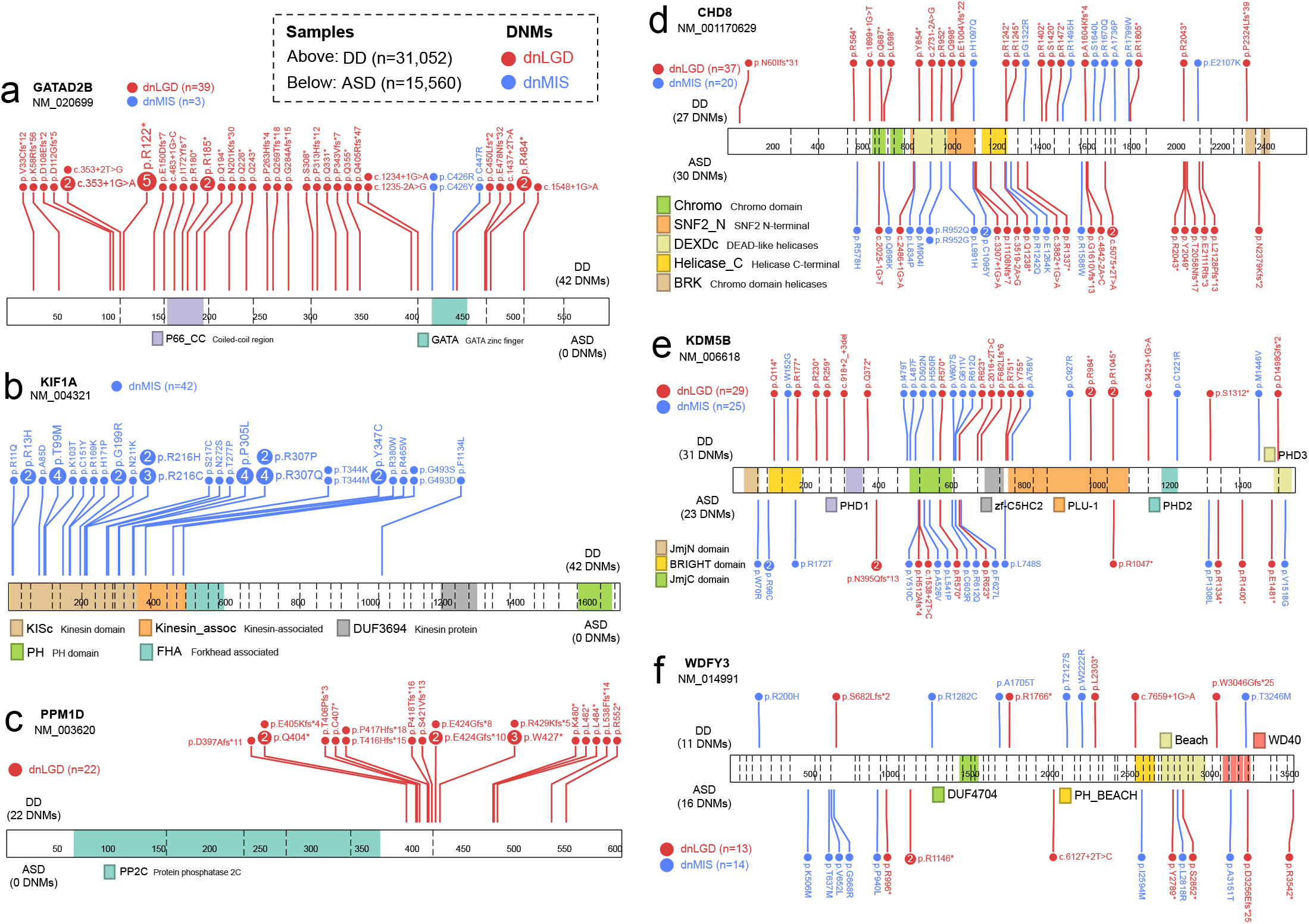
Example genes with mutation class and phenotype-specific DNM pattern. Linear protein diagrams are present with size and exons split by vertical dash lines. Domains are indicated in color blocks with a short description, the total number of dnLGD (red) and dnMIS (blue) variants for each gene was also provided. Recurrent DNMs are indicated by larger circles with the number of recurrences inside. Number of samples plotted: ASD (n = 15,560), DD (n = 31,052). **(a)** *GATAD2B* with DNMs exclusively in DD patients and also enriched for dnLGD variants. **(b)** *KIF1A* only has dnMIS variants and are exclusively in DD patients. **(c)** *PPM1D* only has dnLGD variants and are exclusively in DD patients. **(d)** *CHD8*, **(e)** *KDM5B*, and **(f)** *WDYF3* all have dnLGD and dnMIS variants in both DD and ASD patients, although no phenotype-specific significance, but tends to have more DNMs in ASD than in DD patients when considering the sample size.

### Genes with biased burden by mutation class

In the combined NDD group (n = 46,612), the majority of genes (92.7%, 10,578/11,409) show a greater number of dnMIS when compared to dnLGD variants, which is consistent with expectations. However, a subset of 7.3% (831/11,409) show the reverse pattern with more dnLGD than dnMIS variants (e.g., *ARID1B, ADNP, KMT2A, DYRK1A, CTNNB1, MED13L, POGZ, SETD5, GATAD2B*, etc.), suggesting different mechanisms of pathogenicity or phenotypic manifestation by mutational class. A directly comparison of dnLGD and dnMIS variant count of each gene in the combined NDD cohort (n = 11,409 genes), we identify 12 genes (*ARID1B, ADNP, KMT2A, DYRK1A, CTNNB1, SETD5, GATAD2B, WAC, ASXL1, ASXL3, ARID2*, and *PPM1D*) significantly enriched for dnLGD when compared to dnMIS variants (two-sided Fisher’s exact test, FDR 5%). By contrast, there are 41 genes with significantly greater burden of dnMIS over dnLGD variants (e.g., *DYNC1H1, GRIN2B, CHD3, CACNA1A, SCN8A*, etc.) (Figure 3, Table S11). Among the above genes with a bias by mutation class, *PPM1D* shows exclusively dnLGD variants (n = 22), while 23 genes show only dnMIS variants (e.g., *KIF1A, KCNQ2, PTPN11, SMARCA2*, etc.) in this dataset (Figure 4).

### Sex-based DNM enrichment and burden analysis

The male-biased predominance is well-established for NDDs especially among ASD, but the underlying mechanism remains largely unknown. We further investigated this sex bias in a subset of probands (19,319 males and 8,132 females) where the sex information of the proband is available (Table 1). Using the three models and treating males and females as two distinct groups, we identified 119 and 90 candidate genes with potential male or female bias for DNM, respectively (union FDR 5%) (Tables S12-S13). After applying the most stringent exome-wide significance criteria (intersection FWER 5%), five male-enriched (*ASH1L, ARID2, MBD5, NSD1*, and *SIN3A*) and 17 female-enriched genes (*DDX3X, MECP2, SMC1A, WDR45, HDAC8, NEXMIF, USP9X, CHD7, ZC4H2, CASK, KAT6B, KIF1A, MEF2C, NAA15, PABPC1, PPP2R1A*, and *HNRNPU*) were identified (Figure 3d). Directly comparing of the DNM counts in males versus females, we also identify five genes (*DDX3X, MECP2, SMC1A, WDR45*, and *HDAC8*) with a significant excess of DNM in females (two-sided Fisher’s exact test, FDR 5%) (Table 3, Figure 5). Notably, those five genes all map to the X chromosome with either no DNMs or a single DNM observed in males suggesting that such mutations may be lethal during male embryonic development. Although no individual gene shows a significant male excess when comparing DNM counts between the sexes, several genes show interesting trends. For example, *ASH1L* (aka *KMT2H*), for example, is a histone-lysine N-methyltransferase enzyme with 22 DNMs in males versus only two DNMs observed in females; *KMT2E* encodes a member of the lysine N-methyltransferase-2 family of chromatin modifiers, is associated with O’Donnell-Luria-Rodan syndrome, and shows a male bias in our data (eight DNMs in males verse none in females); finally, *KDM6B* shows eight DNMs in males compared to a single event in females and encodes a histone H3K27 demethylase that specifically demethylates di- or trimethylated lysine-27 (K27) of histone H3 (Table 3, Figure 5).

**Table 3.** Top genes with sex-specific DNM enrichment.

| Gene | Chr | Female DNMs<br>(dnLGD/dnMIS) | Male DNMs<br>(dnLGD/dnMIS) | minimum<br>p-value | minimum<br>q-value | FDR 5%<br>significance | FWER 5%<br>significance |
| --- | --- | --- | --- | --- | --- | --- | --- |
| Top 10 genes with female-specific DNM enrichment (n = 8,132) |  |  |  |  |  |  |  |
| **DDX3X** | X | 52 (24/28) | 0 (0/0) | 4.24E-57 | 4.16E-53 | All | All |
| **MECP2** | X | 22 (5/17) | 1 (1/0) | 2.8E-27 | 2.65E-23 | All | All |
| **SMC1A** | X | 13 (8/5) | 0 (0/0) | 8.88E-15 | 1.2E-11 | All | All |
| **WDR45** | X | 12 (10/2) | 1 (0/1) | 4.72E-26 | 8.13E-23 | All | All |
| **HDAC8** | X | 11 (5/6) | 0 (0/0) | 7.05E-12 | 3.68E-09 | All | All |
| USP9X | X | 12 (9/3) | 3 (0/3) | 8.34E-14 | 1.02E-10 | All | All |
| CASK | X | 9 (7/2) | 2 (2/0) | 1.94E-14 | 2.45E-11 | All | All |
| KIAA2022 | X | 9 (8/1) | 2 (2/0) | 8.1E-14 | 5.43E-11 | All | All |
| ZC4H2 | X | 6 (6/0) | 0 (0/0) | 1.66E-14 | 2.33E-11 | All | All |
| HNRNPU | 1 | 7 (7/0) | 3 (1/2) | 5.58E-40 | 2.64E-36 | All | All |
| Top 10 genes with male-specific DNM enrichment (n = 19,319) |  |  |  |  |  |  |  |
| ASH1L | 1 | 2 (2/0) | 22 (18/4) | 6.72E-26 | 1.59E-22 | All | All |
| ARID2 | 12 | 2 (1/1) | 11 (10/1) | 2.23E-14 | 2.3E-11 | All | All |
| MBD5 | 2 | 2 (2/0) | 10 (6/4) | 5.86E-09 | 2.89E-06 | All | All |
| WDFY3 | 4 | 6 (1/5) | 14 (9/5) | 3.09E-09 | 1.72E-06 | All | CH&DR |
| KCNQ3 | 8 | 1 (0/1) | 8 (0/8) | 8.27E-10 | 4.31E-07 | All | CH&DNW |
| CHAMP1 | 13 | 2 (2/0) | 7 (5/2) | 1.91E-09 | 1.1E-06 | All | DR&DNW |
| ANK2 | 4 | 3 (2/1) | 15 (10/5) | 8.99E-10 | 5.69E-07 | All | DR |
| TNRC6B | 22 | 1 (0/1) | 8 (4/4) | 5.88E-05 | 0.017217 | All | - |
| KDM6B | 17 | 1 (1/0) | 8 (4/4) | 1.54E-05 | 0.004873 | DR&DNW | - |
| KMT2E | 7 | 0 (0/0) | 8 (3/5) | 0.000131 | 0.018551 | DNW | - |

**Figure 5.**
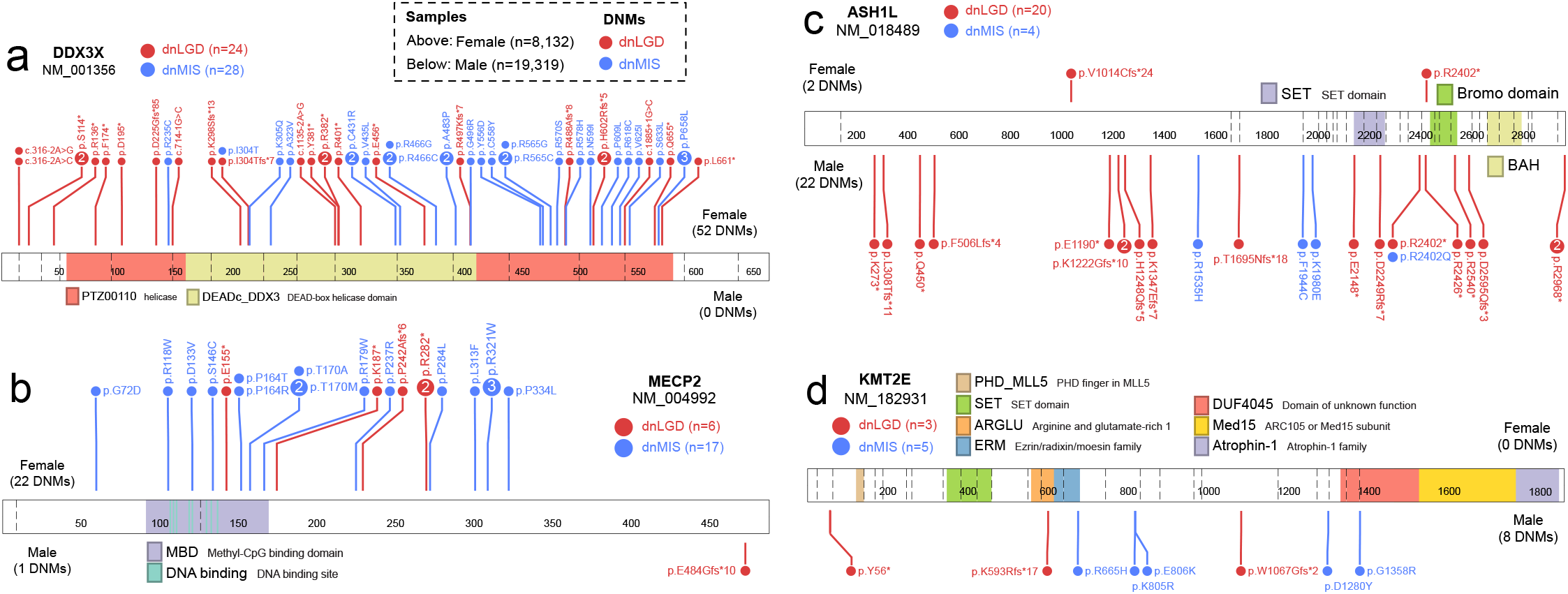
Example genes with sex-biased DNM pattern. Linear protein diagrams are plotted in same way as in Figure 4. Number of samples plotted: female (n = 8,132) and male (n = 19,319). **(a)** *DDX3X* and **(b)** *MECP2* DNMs are almost exclusively in females and with female-only DNM burden and enrichment significance. **(c)** *ASH1L* and **(d)** *KMT2E*, while no sex-bias DNM burden significance, but show male-specific DNM enrichment significance and more DNMs in males than females.

### Functional and neuronal properties of NDD genes

We focus on investigating the functional properties of the highest confidence gene set (HC138) by considering both PPI as well as gene expression trends and differences in the adult and developing human brain. A PPI network analysis using STRING highlights five distinct network clusters (p < 1e-16) (Figure 6a). These include previously described networks such as chromatin binding (GO: 0003682, p = 2.25e-22), ion channel (GO: 0022839, p = 8.43e-13), lysine degradation (hsa00310, KEGG, p = 6.86e-09), and the GABAergic synapse (hsa04727, KEGG, p = 8.78e-08). We identify 20 top “hub” genes, where we observe an excess of PPI interactions supported by half or more of the 12 models in CytoHubba^36^ (Table S14). Tissue-specific expression analysis (TSEA), as expected, shows enrichment in the brain (p = 0.003), with cell-type-specific expression analysis (CSEA) further highlighting expression in specific brain regions from early to late-mid fetal developmental stages; namely: early-mid (p = 1.11e-04) and late-mid fetal (p = 0.005) amygdala, early fetal cerebellum (p = 5.4e-04), early to late fetal cortex (p = 4.87e-06), early to early-mid fetal striatum (p = 1.97e-04), and early fetal thalamus (p = 0.002). Overall, the HC138 genes are more highly expressed prenatally, consistent with the early developmental impact of variation in these genes and NDD (Figure S6).

**Figure 6.**
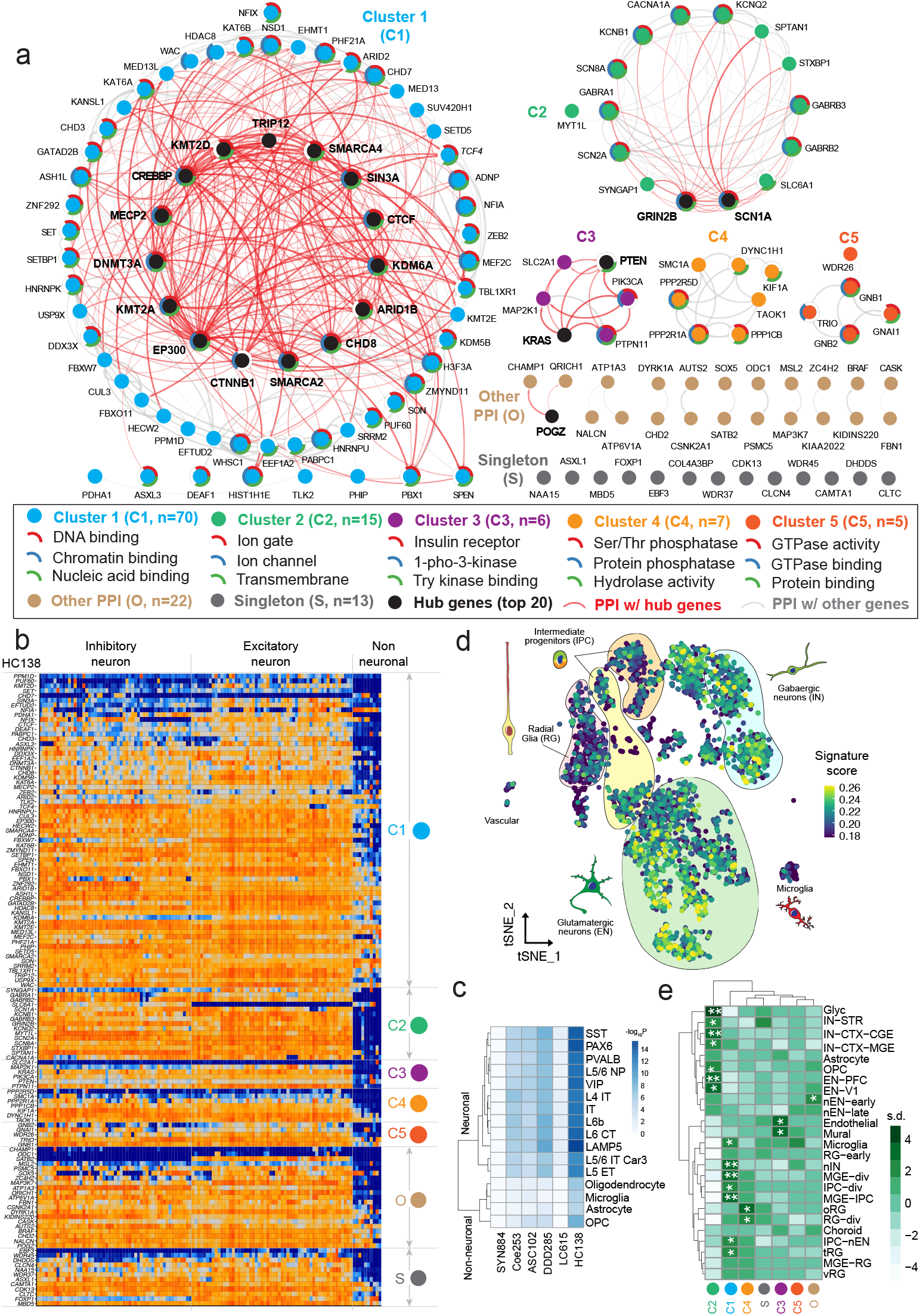
PPI analysis and pan-neuronal expression of the highest confidence genes. **(a)** PPI analysis identified five main clusters (C1-C5), as well as 22 genes in another smaller PPI group (O) and 13 singleton genes (S) using STRING for the HC138 genes with the highest confidence. The top three GO functions were indicated by pie chart in color (top 1 in red, top 2 in blue, and top 3 in green), if applicable, outside each gene dot with a short name in the legend. We also defined 20 top hub genes (black dot and bolded name) supported by at least half of 12 statistical methods in CytoHubba^14^. The interacted edges with hub genes are shown in red curves with the rest in grey curves. The degree of the color and the width of the curve indicate the degree of the interaction. **(b)** Expression heatmap of the HC138 genes in 120 cell types identified across six human neocortical areas grouped by PPI clusters with higher expression (orange) and lower expression (blue). The cell types were grouped by transcriptomic similarity, and the major branches correspond to inhibitory and excitatory neurons and non-neuronal cells. Labels on right side indicate clusters based on PPI analysis, and gene names on the left side. **(c)** Heatmap of -log10-transformed p-values (Bonferroni-corrected for multiple testing) of a Kolmogorov–Smirnov test for the difference in the expression levels of each gene set (rows) in each cell subtype (columns, full name described in Methods) compared to a control set of genes (HC138 and LC615 are the gene sets with the highest and lowest confidence identified in this study; DDD285^16^, ASC102^15^, and Coe253^17^ are significant genes reported previously; SYN884, the control set, includes 844 genes with dnSYN [n > 2] in the 46,612 NDD samples in this study). **(d)** Gene signature score of the HC138 genes computed per cell. Briefly, IN – interneurons, EN excitatory neurons, IPC – intermediate progenitor cells, MGE – medial ganglionic eminence, CGE – caudal ganglionic eminence, OPC – oligeodendrocyte progenitor cells, tRG – truncated radial glia, oRG – outer radial glia, vRG – ventral radial glia, CTX – cortex, V1 – visual cortex, PFC – prefrontal cortex, STR striatum. **(e)** The heatmap shows the standard deviation (s.d.) from the mean expression value of each cluster of genes. Positive values are upregulated compare to the mean, and negative values are downregulated compare to the mean. Significance is derived from bootstrapping and labeled with asterisk (* means p < 0.05, ** means FDR p < 0.05 after Benjamini–Hochberg correction). The HC138 genes were used as the background gene set.

We also explored the enrichment of NDD genes (ASC102^15^, Coe253^17^, DDD285^16^, and LC615 and HC138 in this study) at cellular resolution from expression datasets obtained from the developing and adult human cerebral cortex^37^. Overall, the NDD genes show broad expression across adult human cortical cell types, including inhibitory and excitatory neurons and, to a lesser extent, non-neuronal cells (Figure 6b). All PPI-defined clusters show a similar pattern of pan-neuronal enrichment. While the signal is less pronounced for the LC615 genes, it is still qualitatively more than the 447 genes with non-synonymous DNMs (n > 1) in siblings (Figures S7a,b). We tested for expression enrichment in neuronal and non-neuronal cells for NDD versus control gene sets by comparing expression levels for cell subtypes (Figure 6c, Figure S7c) and the number of clusters that express genes for broad cell classes (Figure S8). All NDD gene sets are significantly enriched in neuronal subtypes, and all gene sets (except the LC615 genes) are enriched in non-neuronal types (Bonferroni-corrected p < 0.05, Kolmogorov–Smirnov test). While different neuronal subtypes show similar enrichments, we note that oligodendrocyte progenitor cells show the greatest enrichment when compared to other non-neuronal cells (Figure 6c). If we use the pan-neuronal expression signature as a further classifier of functional enrichment, we identify 194 of the LC615 gene set meet these criteria as are pan-neuronal genes (Table S8).

To further dissect the neuronal signature, we investigated single-nucleus transcriptomic datasets from the developing human cerebral cortex. At the single-cell level, we find, once again, that HC138 genes are also broadly enriched across inhibitory and excitatory neurons (Figures 6d, e). To test which neuronal subtype shows greater specificity, we performed conditional enrichment analysis between the neuronal lineage populations, and found excitatory neurons to have an independent signal with respect to controlling interneuron signal, while neither enrichment nor significance (p = 1 for all) is reached in any neuronal lineage population when controlling for excitatory neurons (Figure S9). Among excitatory neuron clusters, we find the HC138 genes are most strongly enriched in visual cortex neurons, which is consistent with recently reported changes in gene expression in the postmortem ASD brain tissue samples^38^. When stratified by PPI-defined cluster membership, we find that cluster 1, which harbors many proteins involved in chromatin modification and gene expression regulation, are enriched in progenitor cells and particularly ventral telencephalic neural progenitor cells (‘MGE-IPC’, ‘MGE-div’) and newborn GABAergic neurons (‘nIN’). Cluster 2, which includes proteins involved in neuronal communication and ion channel function, are enriched in cortical excitatory and inhibitory neurons, including excitatory neurons of the prefrontal cortex (‘EN-PFC’), as previously reported^39^, and caudal ganglionic eminence derived GABAergic neurons (‘IN-CTX-CGE’). Additional analysis of cluster 3 and cluster 4 revealed potential trends but did not reach statistical significance after multiple test correction. Genes in cluster 4 (serine/protease phosphorylation), for example, are biased in their expression towards outer radial glia, which are primate-enriched neural stem cells believed to be responsible for the majority of upper cortical layer neurogenesis in humans^40-42^, while genes in cluster 3 (insulin receptor function) are enriched in vascular endothelial and mural cells. We performed a similar single-nucleus transcriptomic analysis for the top 10 genes showing an enrichment trend for DNM in either males or females. Using the HC138 genes as the background gene set, we observe an enrichment signal for intermediate progenitor cells for the female-enriched genes and no signal for male-enriched genes (Figure S10).

## DISCUSSION

Here we present a comprehensive DNM analysis from 15,560 ASD (6,557 are new) and 31,052 DD patients as well as 5,241 unaffected siblings (3,034 are new) using three different statistical models to increase power to identify risk genes and highlight primary diagnosis, sex, mutation class, and model-specific differences. In the combined NDD group, we identify 615 NDD candidate genes with nominal significance supported by any one or more of the three models (LC615, union FDR 5%), of which, 138 genes reaching exome-wide significance in all three models are defined as the most stringent set (HC138, intersection FWER 5%). Among the LC615 genes, 119 were additionally identified only in the combined NDD, which are not observed when ASD and DD are considered independently (Figure 2c). For example, *SYNCRIP* (OMIM: 616686), a gene already strongly implicated in NDD^43,44^, shows two dnLGD and one dnMIS variants in DD patients as well as two dnLGD variants in ASD patients in this dataset, it reaches FDR significance supported by all three models and FWER significance by CH model when combined but no significance when ASD and DD are analyzed separately. Among the high-confidence genes (HC138), there are 10 genes (*MYT1L, PHF21A, FBN1, PBX1, GNB2, PABPC1, CLCN4, NALCN, PSMC5*, and *CERT1*) reaching genome-wide significance (intersection FWER 5%) only in the combined NDD group, which representing good candidates for further investigation (Figure 2c).

Among the LC615 genes, there are 189 genes with potentially novel associations. We note, however, that 84.1% (159/189) of those novel associations only reach nominal significance (FDR 5%) and thus only represent as potential candidates for future investigation as opposed to genes where DNM pathogenicity has been established. Among the HC138 genes supported by all three models (intersection FWER 5%), we highlight three genes (*MED13, NALCN*, and *PABPC1*) reaching exome-wide significance first in this study. *MED13*, a gene that encodes a component of the CDK8-kinase module, has previously been identified through case reports and its phenotype investigated as part of GeneMatcher collaboration^30^. Although *NALCN* was not previously reported as significant in large WES projects of DNM in ASD or DD separately, the gene now reaches the most stringent significance in this study by combining ASD and DD patients. In addition, Chong and colleagues^31^ recently described an autosomal dominant disorder associated with DD and multiple congenital contractures of the face and limbs, extending the phenotype beyond Freeman-Sheldon syndrome to include the impaired cognitive phenotype. Another gene, *PABPC1* encodes a poly-A RNA binding protein and is thought to play a critical role in the structure of the translation initiation complex^45^. We identify 15 dnMIS variants in ASD and DD patients and the gene, to our knowledge, reaches genome-wide significance for an excess of DNM for the first time.

As a whole, the genes identified in this study show a consistent pattern of pan-neuronal expression, which appear to be driven in large part by excitatory neurons. A comparison of single-nucleus transcriptomic data with specific functional PPI networks reveals a more nuanced relationship where specific functionally related genes clearly associate with specific cellular lineages in the brain. Four of the five PPI networks in this study show differential enrichment in specific lineages ranging from rapidly dividing progenitor cells (cluster 1) and outer radial glial cells (cluster 4) to established excitatory and inhibitory neuronal lineages (cluster 2) to cell types associated with vascularization and maintenance of blood-brain barrier (cluster 3). Although not all of these findings are yet statistically significant and will become refined as new highly curated cellular-resolution datasets become available, they suggest possible contribution of these specific cell types to a subset of NDD disease phenotypes and are consistent with observations from studies of large effect size copy number variations on cortical development^46,47^. If functional networks and definition of cell types and circuits are important targets of future therapeutics, distinguishing patients with DNM in these specific genes and further exploration of these PPI networks will be important areas of future research.

Given the locus heterogeneity of ASD and DD, it will be important to continue to expand WES and WGS of parent–child trios in order to identify all the genes associated with excess DNM in NDD. The novel candidate genes identified here with different levels of significance provide an important resource for further genetic and functional characterization. Severe dnMIS30 or dnLGD mutations among the HC138 and LC615 genes account for only 4,667 (10.0%) and 2,872 (6.2%) of the NDD families in this cohort, respectively. Thus, a large number of risk genes are awaiting discovery. Our analyses highlight the value of combining ASD and DD data in increasing power to identify NDD risk genes. While there is ample evidence of DD-specific genes, we find little or no support for ASD-specific genes although do identify candidates with trends toward ASD diagnostic enrichment but none reach statistical significance yet (e.g., *CHD8, KDM5B, WDFY3*)^20^.

However, there are also several limitations and caveats with this study. The size of the DD cohort was double to that of the ASD patients and, as a result, favored the discovery of DD-specific genes where the yield of DNM is greater. The Simons Foundation Powering Autism Research for Knowledge (SPARK) project and other collaborative initiatives aim to generate WES data from over 50,000 ASD families over the next few years and will help rectify this imbalance^48^. Another limitation of this meta-analysis is that not all underlying sequence data or phenotypic information was available or could be accessed for all published cohorts. There are only ∼50% of the raw sequencing data could be reprocessed using the exact same parameters to create a harmonized dataset. Moreover, even basic information regarding patient sex or the ability to match DNM calls to specific samples was limited to some published cohorts. Such impediments need to be rectified for the benefit of the research community and the families who contributed their DNA for the purpose of research. Finally, most of the cohorts included in this study originate from WES whose platform, quality, and coverage varies more widely than WGS data. It has been estimated that 5-8% of genic regions are inaccessible by earlier WES platforms^49^. In the near future, repeating these analyses with WGS data or even ultimately generated by long-read sequencing technologies will help identify new risk genes or loci where may miss by exome sequencing.

## ACKNOWLEDGEMENTS

We are grateful to all the families who participated in the study. We thank Tonia Brown for assistance in editing this manuscript. The DDD study presents independent research commissioned by the Health Innovation Challenge Fund (grant number HICF-1009-003), a parallel funding partnership between the Wellcome Trust and the Department of Health, and the Wellcome Trust Sanger Institute (grant number WT098051). The views expressed in this publication are those of the authors and not necessarily those of the Wellcome Trust or the Department of Health. The study has UK Research Ethics Committee approval (10/H0305/83, granted by the Cambridge South REC, and GEN/284/12 granted by the Republic of Ireland REC). The research team acknowledges the support of the National Institute for Health Research, through the Comprehensive Clinical Research Network. This work was supported, in part, by US National Institutes of Health (NIH) grants (R01MH101221 and U01MH119705) and a grant from the Simons Foundation (SFARI #608045) to E.E.E.; E.E.E. is an investigator of the Howard Hughes Medical Institute.

## AUTHOR CONTRIBUTIONS

T.W. and E.E.E. designed the study; T.W. analyzed, collected and harmonized the dataset, and performed the data analyses. C.K., T.E.B., and T.J.N. helped with the gene expression analyses. M.A.G. contributed in interpretation of the candidate genes. B.H. and Y.M. helped with figure preparations. C.G. contributed some of the underlying data of RUMC cohort. T.W. and E.E.E. wrote the manuscript with input from other authors. All authors reviewed and approved the manuscript.

## METHODS

### Cohorts and samples

This study was approved by the University of Washington Institutional Review Board #STUDY00000383, Genetics Consortium Repository. Informed consent was obtained from all subjects by each of the corresponding study cohort. We collected eight WES and three WGS parent–child cohorts (>100 trios), including over 44,800 families from both published and publicly available data (Table S1). We preferentially retain WGS over WES data for cohorts that have both types of data available. For example, we only included the Centers for Common Disease Genomics (CCDG) genomes for Simons Simplex Collection (SSC) samples^26^. For cohorts with continuous publications, like for the Autism Sequencing Consortium (ASC), Deciphering Developmental Disorders (DDD), and Radboud University Medical Center (RUMC) samples, we only included the final samples from their latest study with potential sample duplicates removed. We also excluded any potential overlaps in the literature; for example, we excluded all SSC samples used in the ASC paper^15^ for potential redundancy with the CCDG SSC genomes. After all those control measures, we also ran KING^50^ (v1.4) for samples with the underlying sequencing data available for further detection of potential sample overlap. KING uses identical by state (IBS) to estimate pairwise relatedness between samples; any samples with a kinship value >0.35 were considered as potential sample duplicates. We first checked if the potential duplicates were known as monozygotic twin pairs or known duplicates within a cohort (some individuals had both blood and cell line DNA sequenced for quality control purposes). Only one sample was retained from each duplicate pair in downstream analyses—blood DNA was preferred if both blood and cell line sequencing data were available.

### DNM discovery and integration

DNMs were identified by analyzing/reanalyzing the underlying sequencing data wherever available for the five cohorts using the same pipeline. Specifically, DNMs were harmonized by reanalyzing 70,172 samples (46.5%), including 24,520 families within two categories. First, for ASD cohorts with genome sequencing data from the CCDG study, including SSC and the Study of Autism Genetics Exploration (SAGE), raw single-nucleotide variant/insertion or deletion (SNV/indel) variants were called (on hg38) independently using four different callers: GATK^51^, FreeBayes^52^, Platypus^53^, and Strelka2^54^. Downstream DNM discovery was based on genotype, which is required only if the offspring has the alternative allele (with genotype as 0/1 or 1/1) but is not observed in either of the parents (with genotype as 0/0). Candidate DNMs needed to have the support of at least two of the four callers; and then variants from Platypus with a filter of LowGQX or NoPassedVariantGTs were removed, and Strelka2 variants had to have the filter field equal to PASS. For variants on the X chromosome, we separately considered variants in the pseudoautosomal regions (chrX:10000-2781479, chrX:155701382-156030895, hg38) and the X/Y duplicative transposed region (chrX:89201803-93120510, hg38). Candidate DNMs were then converted to hg19 for downstream integration. Second, for part of other WES cohorts, including SPARK_pilot, SPARK_WES_1, and DDD, raw SNV/indel variants were called (on hg19) independently using GATK and FreeBayes. Downstream DNM discovery was based on genotype as applied in genome cohorts. Candidate DNMs needed to have the support from both callers, which is the intersection set by GATK and FreeBayes. Beyond the above measures, we also applied the following variant-level filters: allele balance (AB = 0 in both parents, and AB > 0.25 in the child), read depth (DP > 9 for all family members), child genotype quality (GQ > 20 by both GATK and FreeBayes). For the rest cohorts included in studies with no underlying sequencing data available, DNMs were collected from each corresponding publication. DNMs on hg38 were first converted to hg19; the final integrated DNMs were all on hg19. To combine DNMs between WES and WGS datasets, we restricted DNMs to a well-covered coding region^55^ (average DP > 20X) generated by accessing the WES data from the SSC and DDD study. We also removed all DNMs in the segmental duplication regions, recent repeat and low-complexity regions, or centromeric and telomeric regions. We excluded variants in a homopolymer A or T of length 10 or greater, and the variants with a reference or alternative allele with greater than 10 bp, to remove potential sequencing errors. Beyond all the above filtering and sample duplicate exclusions, we further excluded samples with more than 10 coding DNMs as outliers and removed specific DNMs that were observed in more than five different unrelated individuals for frequency control. All of the above strict measures yielded a total of 46,612 nonredundant NDD cases with a primary diagnosis of ASD (n = 15,560) or DD (n = 31,052), and also unaffected siblings (n = 5,241) in the integrated *de novo* enrichment analysis (Table S1). To ensure uniformity, the same version of CADD score (v1.3) and VEP annotation (Ensembl GRCh37 release 94) were applied, and the analysis was restricted to the canonical transcript with the most deleterious annotation.

### Statistical analyses

*De novo* enrichment analyses were performed independently for ASD, DD, and NDD samples by using three statistical models: the CH model, denovolyzeR, and DeNovoWEST. All three methods apply their own underlying variant rate estimates (denovolyzeR and DeNovoWEST use the same rate while CH model is different) to generate the prior probabilities for observing a specific number and class of mutations for a given gene. Briefly, the CH model estimates the number of expected DNMs by incorporating locus-specific transition, transversion, and indel rates and chimpanzee–human coding sequence divergence and the gene length; while denovolyzeR estimates mutation rates based on trinucleotide context and incorporates exome depth, mutational biases such as CpG hotspots, and divergence adjustments based on macaque–human comparisons. DeNovoWEST is a further development beyond denovolyzeR, which scores all classes of coding variants on a unified severity scale based on the empirically estimated positive predictive value of being pathogenic, and incorporates a gene-based weighting derived from the deficit of protein-truncating variants in the general population, further combining missense enrichment by a clustering test. Default parameters were used for all three methods with some minor adjustments, such as in the process of weight creation in DeNovoWEST, fewer numbers of CADD bins for missense and nonsense variants were used for ASD samples (three bins), versus in the DD and combined NDD group where seven bins were used for both, due to the sample size differences as suggested^6^. The expected mutation rate of 1.8 DNMs per exome was set to the CH model as an upper bound baseline. Siblings were also analyzed similarly using the CH model and denovolyzeR, but not run for DeNovoWEST due to the small sample size. We applied two metrics of significance with the union and intersection of three models: first is the FDR significance, the significance threshold (q < 0.05) was corrected exome-wide using Benjamini–Hochberg method by accounting for the total number of genes in each model (18,946 genes in CH model, 19,618 genes in denovolyzeR, and 18,762 genes in DeNovoWEST); the second is a more stringent FWER significance, for which we applied exome-wide Bonferroni multiple-testing correction considering both the largest number of genes among three models (n = 19,618) and the total of tests per gene across the three models. For probands, FWER 5% significance threshold (p < 3.64e-07) was corrected by the Bonferroni method for 19,618 genes and seven tests in the analysis (dnLGD, dnMIS, and dnMIS30 in CH model, dnLGD and dnMIS variants in denovolyzeR, and dnLGD and dnMIS variants in DeNovoWEST). For siblings, FWER 5% significance threshold (p < 5.09e-07) was corrected for 19,618 genes but only five tests in the CH model and denovolyzeR. We excluded genes that show any significance in the siblings from the counting of significant genes in probands (ASD, DD, and NDD). For each variant category, we required each gene to have more than two DNMs to be considered as significant. All statistics were calculated using R (versions 3.6.2).

### Enrichment analyses in recalled and no-recall subsets

We also performed same enrichment analyses using three models in parallel for those two subsets. We identified 323 FDR (132 FWER) significant genes in the recalled subset and 389 FDR (174 FWER) significant genes in the no-recall subset (Tables S15-S16). For those FDR-significant genes, of which 87.3% (282/323) in the recalled subset and 90.0% (350/389) in the no-recall subset overlap with the LC615 genes in combined NDD group (Figure S11), suggesting consistent results in both subsets after data harmonization. However, there are 12 exclusively significant genes in the no-recall subset and one exclusively significant gene (*PABPC1*) in the recalled subset among the HC138 genes in combined NDD group. A closer look found those 12 genes were also reported as significant genes in a recent study^16^, and the significant signal was driven by DNMs almost exclusively from GeneDx, which the raw data is not available for recalling. For example, all DNMs in three genes (*ZEB2, PDHA1*, and *SLC2A1*) are from GeneDx probands (n = 18,783) and none among the other five cohorts (n = 8,454) in the no-recall subset (Figure S11). This is consistent with the original study where the majority of the DNMs are from GeneDx cohort, with very few from the DDD and RUMC samples. This draws attention to the significance of such genes, where the significance signal is mostly driven by DNMs from a single cohort and no underlying data is available for reanalyzing as more quality control.

### PPI analyses and hub genes

PPI network was assessed by searching Multiple Proteins by Names using the online STRING database with default settings. The interaction result was exported as a TSV file and then imported into Cytoscape software for downstream analysis. We used CytoHubba to identify top hub genes (most interacted genes). CytoHubba provides the analyzed results computed by 12 methods, including Degree, clustering coefficient, Edge Percolated Component (EPC), Maximum Neighborhood Component (MNC), Density of Maximum Neighborhood Component (DMNC), Maximal Clique Centrality (MCC), and centralities based on shortest paths, such as Bottleneck (BN), EcCentricity, Closeness, Radiality, Betweenness, and Stress, as previously described^36^. The top 20 genes were supported by the most, and at least half, of the models as top hub genes. The PPI clusters were identified by the Markov Cluster Algorithm (MCL, https://micans.org/mcl/). The top three GO functions were selected from rank order of the functional enrichment from STRING database with default settings.

### Single-nucleus RNA expression analysis

The dataset includes single-nucleus transcriptomes from 49,495 nuclei across multiple human cortical areas. Individual cortex layers were dissected from tissues covering the middle temporal gyrus (<u>MTG</u>), anterior cingulate cortex (<u>ACC</u>; also known as the ventral division of medial prefrontal cortex, A24), primary visual cortex (<u>V1C</u>), primary motor cortex (<u>M1C</u>), primary somatosensory cortex (<u>S1C</u>), and primary auditory cortex (<u>A1C</u>) derived from human brain. Nuclei were dissociated and sorted using the neuronal marker NeuN. Nuclei were sampled from postmortem and neurosurgical (MTG only) donor brains and expression was profiled with SMART-Seq v4 RNA-sequencing. The data are available from the Allen Institute for Brain Science website for analysis (https://celltypes.brain-map.org/rnaseq/human_ctx_smart-seq) and download (https://portal.brain-map.org/atlases-and-data/rnaseq/human-multiple-cortical-areas-smart-seq). Unsupervised clustering with Seurat identified 120 distinct transcriptomic clusters, including 54 GABAergic (inhibitory) neuronal, 56 glutamatergic (excitatory) neuronal, and 10 non-neuronal cell types. Heatmaps were constructed of log-normalized trimmed mean expression (excluding the 25% lowest and 25% highest expression values), log_2_(CPM + 1), of NDD and control gene sets across cell types. Genes were ordered by the number of cell types with trimmed mean expression > 1. For each cell class (GABAergic and glutamatergic neurons and non-neuronal cells), the number of cell types with trimmed mean expression > 1 for NDD risk genes and control genes were quantified and visualized as empirical cumulative distributions (Figure S8). A Kolmogorov–Smirnov test was used to reject the null hypothesis that the cell type count distributions were the same between each gene set and the control DNM gene set. P-values were Bonferroni-corrected for multiple testing. Similarly, for each cell subclass (e.g., SST interneurons or L6b excitatory neurons), the trimmed mean expression levels of NDD risk genes and control genes were quantified and visualized as empirical cumulative distributions (for example, Figure S7c). A Kolmogorov– Smirnov test was used to reject the null hypothesis that the expression distributions were the same between each gene set and the control DNM gene set. P-values were Bonferroni corrected for multiple testing and -log_10_-transformed and visualized as a heatmap with columns corresponding to cell subclasses ordered by Ward’s clustering and rows corresponding to gene sets. Full names and detailed descriptions of each cell subtype in the heatmap (Figure 6c) are: GABAergic interneuron (LAMP5: LAMP5 expressing GABAergic neuron, PAX6: PAX6 expressing GABAergic neuron, PVALB: PVALB expressing GABAergic neuron, SST: SST expressing GABAergic neuron; VIP: VIP expressing GABAergic neuron); Glutamatergic neuron (IT: Intratelencephalic neuron, L4 IT: Layer 4 Intratelencephalic neuron, L5 ET: Layer 5 Extratelencephalic neuron, L5/6 IT Car3: Layer 5/6 Intratelencephalic neuron that selectively expresses Car3, L5/6 NP: Layer 5-6 Near-projecting neuron, L6 CT: Layer 6 Corticothalamic neuron, L6b: Layer 6b neuron); and Non-neuronal cell (Astrocyte, Microglia, Oligodendrocyte, OPC: oligodendrocyte progenitor cell).

### Tissue and cell-type-specific expression of significant genes

scRNA-seq data were pulled from http://cells.ucsc.edu/?ds=cortex-dev and CPM counts were quasi normalized unto unique molecular identifies (UMI) using quminorm (https://github.com/willtownes/quminorm). Cells were then regrouped by their broad parent cell types with unknown cell types filtered out. SCTransform (https://github.com/ChristophH/sctransform) was used to normalize the UMI counts from quminorm. The corrected counts from SCTransform were used as input into Expression Weighted Celltype Enrichment (EWCE) following the default parameters with two levels of annotations based on clusters and clusters split by sex. Bootstrapping parameters in EWCE: 10000 repetitions with the LC615 genes as background in the unconditional enrichment and HC138 for the controlled experiments. Cluster-specific analysis within the most stringent gene set and top sex specific genes (10 for female and 10 for male) used the HC138 genes as background. Online TSEA and CSEA tools were used to determine the enrichment of expression across brain regions and cell types^56^. The expression among these tissues was compared using Fisher’s exact tests and followed by Benjamini–Hochberg correction.

### Assessment of gene intolerance scores

To assess a gene’s intolerance to variation, we applied the ExAC-based residual variance to intolerance score (RVIS) and missense constraint scores (mis_Z scores), as well as the gnomAD-based “loss-of-function observed/expected upper bound fraction” (LOEUF). LOEUF score is a conservative estimate of the observed/expected ratio, based on the upper bound of a Poisson-derived confidence interval around the ratio. It ranges from 0 to 2, with lower LOEUF scores indicating stronger selection against predicted loss-of-function variation in a given gene, and a cut-off value is suggested as 0.35. The mis_Z score indicates a gene’s intolerance to missense variants, positive scores indicate more constraint, and negative scores indicate less constraint. A greater Z-score indicates more intolerance to the class of variation. RVIS gene score was based on ExAC v2 release 2.0 (accessed: March 15^th^, 2017). As of this release we use CCDS release 20 and Ensembl release 87 annotations. The score was converted into percentile by ranking all genes from most intolerant to least. For example, percentile of 1% means the gene is amongst the top 1% of the most intolerant genes. Wilcoxon two-sample test was performed in R (versions 3.6.2) with the wilcox.test function.

## SUPPLEMENTAL INFORMATION

Supplemental_Information.pdf

- Figures S1-S11
- Tables S1, S2, S9

Supplemental_Tables.xlsx

- Tables S3-S8, S10-S16

## DATA AND CODE AVAILABILITY

The underlying genomic and phenotypic data in the recalled subset are available from the following resources: the sequencing and phenotype data for the SSC cohort are available to approved researchers at SFARI Base (accession IDs: SFARI_SSC_WGS_p, SFARI_SSC_WGS_1, and SFARI_SSC_WGS_2). WGS data from the SAGE samples is available at dbGaP under accession phs001740.v1.p1. Family-level FreeBayes and GATK VCF files for SSC and SAGE samples are available at dbGaP under accession phs001874.v1.p1 and also at SFARI Base under accession: SFARI_SSC_WGS_2a. The genomic and phenotypic data for the SPARK study is available by request from SFARI Base (accession IDs: SFARI_SPARK_pilot, SFARI_SPARK_WES_1). The genomic and phenotypic data for the DDD study is available by request from the European Genome-phenome Archive (EGA) under study EGAS00001000775. The DNMs in the no-recall subset are retrieved from each of the corresponding publications (Table S2). The final harmonized DNMs, from both the recalled and no-recall subsets, used in the study are available in Table S3. This study did not generate new unique reagents. Code used for analyses and figures are available upon request. Further information and requests for resources should be directed to and will be fulfilled by the Lead Contact, Evan E. Eichler. Software and databases used in this study are publicly available as follows:

denovo-db: http://denovo-db.gs.washington.edu/;

SPARK: https://sparkforautism.org;

Ensembl VEP (GRCh37): http://grch37.ensembl.org/Homo_sapiens/Tools/VEP/;

CADD score: https://cadd.gs.washington.edu/;

FreeBayes: https://github.com/ekg/freebayes;

CH model: https://github.com/tianyunwang/CH-model;

denovolyzeR: https://github.com/jamesware/denovolyzeR;

DeNovoWEST: https://github.com/queenjobo/DeNovoWEST;

quminorm: https://github.com/willtownes/quminorm;

SCTransform: https://github.com/ChristophH/sctransform;

UCSC Cell Browser: https://cells.ucsc.edu;

TSEA tool: http://genetics.wustl.edu/jdlab/tsea/;

CSEA tool: http://genetics.wustl.edu/jdlab/csea-tool-2/;

RVIS score: http://genic-intolerance.org/;

STRING: https://string-db.org/;

CytoScape: https://cytoscape.org/;

CytoHubba: http://apps.cytoscape.org/apps/cytohubba;

ProteinPaint: https://proteinpaint.stjude.org/;

DDG2P: https://www.ebi.ac.uk/gene2phenotype;

SFARI Gene: https://gene.sfari.org/;

SFARI Base: http://base.sfari.org;

EGA: https://ega-archive.org/;

OMIM: https://omim.org;

gnomAD: https://gnomad.broadinstitute.org/.

## ABBREVIATIONS

DNM: *de novo* mutation
NDD: Neurodevelopmental disorder
ASD: Autism spectrum disorder
DD: Developmental disorder
ID: Intellectual disability
WES: Whole-exome sequencing
WGS: Whole-genome sequencing
CCDG: Centers for Common Disease Genomics
SFARI: Simons Foundation Autism Research Initiative
SPARK: Simons Foundation Powering Autism Research for Knowledge
SSC: Simons Simplex Collection
ASC: Autism Sequencing Consortium
DDD: Deciphering Developmental Disorders
RUMC: Radboud University Medical Center
CADD score: combined annotation dependent depletion score
LGD: likely gene-disruptive
dnLGD: *de novo* LGD variant
dnSYN: *de novo* synonymous variant
dnMIS: *de novo* missense variant
dnMIS30: *de novo* missense variant with CADD score greater than 30
FDR: False discovery rate
FWER: Family-wise error rate
PPI: Protein-protein interaction
CSEA: Cell-type-specific expression analysis
TSEA: Tissue-specific expression analysis
DDG2P: Development Disorder Genotype - Phenotype Database
OMIM: Online Mendelian Inheritance in Man
LC615: The 615 genes are with a lower confidence at the union significance (FDR 5%) by one or more of the three models
HC138: The 138 genes are with the highest confidence at the intersection significance (FWER 5%) supported by all three models, which are a subset of the LC615 genes

## The SPARK Consortium

John Acampado ^1^, Andrea J. Ace ^1^, Alpha Amatya ^1^, Irina Astrovskaya ^1^, Asif Bashar ^1^, Elizabeth Brooks ^1^, Martin E. Butler ^1^, Lindsey A. Cartner ^1^, Wubin Chin ^1^, Wendy K. Chung ^1,2^, Amy M. Daniels ^1^, Pamela Feliciano ^1^, Chris Fleisch ^1^, Swami Ganesan ^1^, William Jensen ^1^, Alex E. Lash ^1^, Richard Marini ^1^, Vincent J. Myers ^1^, Eirene O’Connor ^1^, Chris Rigby ^1^, Beverly E. Robertson ^1^, Neelay Shah ^1^, Swapnil Shah ^1^, Emily Singer ^1^, LeeAnne G. Snyder ^1^, Alexandra N. Stephens ^1^, Jennifer Tjernagel ^1^, Brianna M. Vernoia ^1^, Natalia Volfovsky ^1^, Loran Casey White ^1^, Alexander Hsieh ^2^, Yufeng Shen ^2^, Xueya Zhou ^2^, Tychele N. Turner ^3^, Ethan Bahl ^4^, Taylor R. Thomas ^4^, Leo Brueggeman ^4^, Tanner Koomar ^4^, Jacob J. Michaelson ^4^, Brian J. O’Roak ^5^, Rebecca A. Barnard ^5^, Richard A. Gibbs ^6^, Donna Muzny ^6^, Aniko Sabo ^6^, Kelli L. Baalman Ahmed ^6^, Evan E. Eichler ^7^, Matthew Siegel ^8^, Leonard Abbeduto ^9^, David G. Amaral ^9^, Brittani A. Hilscher ^9^, Deana Li ^9^, Kaitlin Smith ^9^, Samantha Thompson ^9^, Charles Albright ^10^, Eric M. Butter ^10^, Sara Eldred ^10^, Nathan Hanna ^10^, Mark Jones ^10^, Daniel Lee Coury ^10^, Jessica Scherr ^10^, Taylor Pifher ^10^, Erin Roby ^10^, Brandy Dennis ^10^, Lorrin Higgins ^10^, Melissa Brown ^10^, Michael Alessandri ^11^, Anibal Gutierrez ^11^, Melissa N. Hale ^11^, Lynette M. Herbert ^11^, Hoa Lam Schneider ^11^, Giancarla David ^11^, Robert D. Annett ^12^, Dustin E. Sarver ^12^, Ivette Arriaga ^13^, Alexies Camba ^13^, Amanda C. Gulsrud ^13^, Monica Haley ^13^, James T. McCracken ^13^, Sophia Sandhu ^13^, Maira Tafolla ^13^, Wha S. Yang ^13^, Laura A. Carpenter ^14^, Catherine C. Bradley ^14^, Frampton Gwynette ^14^, Patricia Manning ^15^, Rebecca Shaffer ^15^, Carrie Thomas ^15^, Raphael A. Bernier ^16^, Emily A. Fox ^16^, Jennifer A. Gerdts ^16^, Micah Pepper ^16^, Theodore Ho ^16^, Daniel Cho ^16^, Joseph Piven ^17^, Holly Lechniak ^18^, Latha V. Soorya ^18^, Rachel Gordon ^18^, Allison Wainer ^18^, Lisa Yeh ^18^, Cesar Ochoa-Lubinoff ^19^, Nicole Russo ^19^, Elizabeth Berry-Kravis ^20^, Stephanie Booker ^21^, Craig A. Erickson ^21^, Lisa M. Prock ^22^, Katherine G. Pawlowski ^22^, Emily T. Matthews ^22^, Stephanie J. Brewster ^22^, Margaret A. Hojlo ^22^, Evi Abada ^22^, Elena Lamarche ^23^, Tianyun Wang ^24^, Shwetha C. Murali ^7^, William T. Harvey ^24^, Hannah E. Kaplan ^25^, Karen L. Pierce ^25^, Lindsey DeMarco ^26^, Susannah Horner ^26^, Juhi Pandey ^26^, Samantha Plate ^26^, Mustafa Sahin ^27^, Katherine D. Riley ^27^, Erin Carmody ^27^, Julia Constantini ^7^, Amy Esler ^28^, Ali Fatemi ^29^, Hanna Hutter ^29^, Rebecca J. Landa ^29^, Alexander P. McKenzie ^29^, Jason Neely ^29^, Vini Singh ^29^, Bonnie Van Metre ^29^, Ericka L. Wodka ^29^, Eric J. Fombonne ^30^, Lark Y. Huang-Storms ^30^, Lillian D. Pacheco ^30^, Sarah A. Mastel ^30^, Leigh A. Coppola ^30^, Sunday Francis ^31^, Andrea Jarrett ^31^, Suma Jacob ^31^, Natasha Lillie ^31^, Jaclyn Gunderson ^31^, Dalia Istephanous ^31^, Laura Simon ^31^, Ori Wasserberg ^31^, Angela L. Rachubinski ^32^, Cordelia R. Rosenberg ^32^, Stephen M. Kanne ^33,34^, Amanda D. Shocklee ^34^, Nicole Takahashi ^34^, Shelby L. Bridwell ^34^, Rebecca L. Klimczac ^34^, Melissa A. Mahurin ^34^, Hannah E. Cotrell ^34^, Cortaiga A. Grant ^34^, Samantha G. Hunter ^34^, Christa Lese Martin ^35^, Cora M. Taylor ^35^, Lauren K. Walsh ^35^, Katherine A. Dent ^35^, Andrew Mason ^36^, Anthony Sziklay ^36^, Christopher J. Smith ^36^

^1^ Simons Foundation, New York, USA

^2^ Columbia University, New York, USA

^3^ Washington University School of Medicine, St. Louis, USA

^4^ University of Iowa Carver College of Medicine, Iowa City, USA

^5^ Oregon Health & Science University, Portland, USA

^6^ Baylor College of Medicine, Houston, USA

^7^ University of Washington School of Medicine & Howard Hughes Medical Institute, Seattle, USA

^8^ Maine Medical Center Research Institute, Portland, USA

^9^ University of California, Davis, Sacramento, USA

^10^ Nationwide Children’s Hospital, Columbus, USA

^11^ University of Miami, Coral Gables, USA

^12^ University of Mississippi Medical Center, Jackson, USA

^13^ University of California, Los Angeles, Los Angeles, USA

^14^ Medical University of Southern Carolina (MUSC), Portland, USA

^15^ Cincinnati Children’s Hospital Medical Center - Research Foundation, Cincinnati, USA

^16^ Seattle Children’s Autism Center/UW, Seattle, USA

^17^ University of North Carolina at Chapel Hill, Chapel Hill, USA

^18^ Department of Child & Adolescent Psychiatry, Rush University Medical Center, Chicago, USA

^19^ Department of Developmental & Behavioral Pediatrics, Rush University Medical Center, Chicago, USA

^20^ Department of Neurological Sciences, Department of Pediatrics, Department of Biochemistry, Rush University Medical Center, Chicago, USA

^21^ Cincinnati Children’s Hospital Medical Center - Research Foundation, Cincinnati, USA

^22^ Boston Children’s Hospital (BCH), Boston, USA

^23^ University of North Carolina at Chapel Hill, Chapel Hill, USA

^24^ University of Washington School of Medicine, Seattle, USA

^25^ University of California, San Diego, School of Medicine, La Jolla, USA

^26^ Children’s Hospital of Philadelphia, Philadelphia, USA

^27^ Boston Children’s Hospital (BCH), Boston, USA

^28^ University of Minnesota, Minneapolis, USA

^29^ Kennedy Krieger Institute, Baltimore, USA

^30^ Oregon Health & Science University, Portland, USA

^31^ University of Minnesota, Minneapolis, USA

^32^ University of Colorado School of Medicine, Aurora, USA

^33^ Department of Health Psychology, University of Missouri, Columbia, USA

^34^ Thompson Center for Autism and Neurodevelopmental Disorders, University of Missouri, Columbia, USA

^35^ Geisinger Autism & Developmental Medicine Institute, Lewisburg, USA

^36^ Southwest Autism Research and Resource Center, Phoenix, USA

## REFERENCES

1. Zablotsky, B. et al. Prevalence and Trends of Developmental Disabilities among Children in the United States: 2009–2017. (2019).

2. First, M.B. Diagnostic and statistical manual of mental disorders, 5th edition, and clinical utility. J Nerv Ment Dis 201, 727–9 (2013).

3. Srivastava, A.K. & Schwartz, C.E. Intellectual disability and autism spectrum disorders: causal genes and molecular mechanisms. Neurosci Biobehav Rev 46 Pt 2, 161–74 (2014).

4. Mefford, H.C., Batshaw, M.L. & Hoffman, E.P. Genomics, intellectual disability, and autism. N Engl J Med 366, 733–43 (2012).

5. Lyall, K. et al. The Changing Epidemiology of Autism Spectrum Disorders. Annu Rev Public Health 38, 81–102 (2017).

6. Zablotsky, B. et al. Prevalence and Trends of Developmental Disabilities among Children in the United States: 2009-2017. Pediatrics 144(2019).

7. Stromme, P. & Diseth, T.H. Prevalence of psychiatric diagnoses in children with mental retardation: data from a population-based study. Dev Med Child Neurol 42, 266–70 (2000).

8. La Malfa, G., Lassi, S., Bertelli, M., Salvini, R. & Placidi, G.F. Autism and intellectual disability: a study of prevalence on a sample of the Italian population. J Intellect Disabil Res 48, 262–7 (2004).

9. Bryson, S.E., Bradley, E.A., Thompson, A. & Wainwright, A. Prevalence of autism among adolescents with intellectual disabilities. Can J Psychiatry 53, 449–59 (2008).

10. de Bildt, A. et al. Interrelationship between Autism Diagnostic Observation Schedule-Generic (ADOS-G), Autism Diagnostic Interview-Revised (ADI-R), and the Diagnostic and Statistical Manual of Mental Disorders (DSM-IV-TR) classification in children and adolescents with mental retardation. J Autism Dev Disord 34, 129–37 (2004).

11. Christensen, D.L. et al. Prevalence and Characteristics of Autism Spectrum Disorder Among Children Aged 4 Years - Early Autism and Developmental Disabilities Monitoring Network, Seven Sites, United States, 2010, 2012, and 2014. MMWR Surveill Summ 68, 1–19 (2019).

12. Matson, J.L. & Shoemaker, M. Intellectual disability and its relationship to autism spectrum disorders. Res Dev Disabil 30, 1107–14 (2009).

13. Iossifov, I. et al. The contribution of de novo coding mutations to autism spectrum disorder. Nature 515, 216–21 (2014).

14. Deciphering Developmental Disorders, S. Prevalence and architecture of de novo mutations in developmental disorders. Nature 542, 433–438 (2017).

15. Satterstrom, F.K. et al. Large-Scale Exome Sequencing Study Implicates Both Developmental and Functional Changes in the Neurobiology of Autism. Cell 180, 568–584 e23 (2020).

16. Kaplanis, J. et al. Evidence for 28 genetic disorders discovered by combining healthcare and research data. Nature (2020).

17. Coe, B.P. et al. Neurodevelopmental disease genes implicated by de novo mutation and copy number variation morbidity. Nature Genetics 51, 106–116 (2019).

18. O’Roak, B.J. et al. Multiplex targeted sequencing identifies recurrently mutated genes in autism spectrum disorders. Science 338, 1619–22 (2012).

19. Ware, J.S., Samocha, K.E., Homsy, J. & Daly, M.J. Interpreting de novo Variation in Human Disease Using denovolyzeR. Curr Protoc Hum Genet 87, 7 25 1–7 25 15 (2015).

20. Myers, S.M. et al. Insufficient Evidence for “Autism-Specific” Genes. Am J Hum Genet 106, 587–595 (2020).

21. Guo, H. et al. Genome sequencing identifies multiple deleterious variants in autism patients with more severe phenotypes. Genet Med 21, 1611–1620 (2019).

22. Feliciano, P. et al. Exome sequencing of 457 autism families recruited online provides evidence for autism risk genes. NPJ Genom Med 4, 19 (2019).

23. Rk, C.Y. et al. Whole genome sequencing resource identifies 18 new candidate genes for autism spectrum disorder. Nat Neurosci 20, 602–611 (2017).

24. Chen, R. et al. Leveraging blood serotonin as an endophenotype to identify de novo and rare variants involved in autism. Mol Autism 8, 14 (2017).

25. Takata, A. et al. Integrative Analyses of De Novo Mutations Provide Deeper Biological Insights into Autism Spectrum Disorder. Cell Rep 22, 734–747 (2018).

26. Wilfert, A.B. et al. Recent ultra-rare inherited mutations identify novel autism candidate risk genes. bioRxiv, 2020.02.10.932327 (2020).

27. Krumm, N. et al. Excess of rare, inherited truncating mutations in autism. Nat Genet 47, 582–8 (2015).

28. Rentzsch, P., Witten, D., Cooper, G.M., Shendure, J. & Kircher, M. CADD: predicting the deleteriousness of variants throughout the human genome. Nucleic Acids Res (2018).

29. Kircher, M. et al. A general framework for estimating the relative pathogenicity of human genetic variants. Nat Genet 46, 310–5 (2014).

30. Snijders Blok, L. et al. De novo mutations in MED13, a component of the Mediator complex, are associated with a novel neurodevelopmental disorder. Hum Genet 137, 375–388 (2018).

31. Chong, J.X. et al. De novo mutations in NALCN cause a syndrome characterized by congenital contractures of the limbs and face, hypotonia, and developmental delay. Am J Hum Genet 96, 462–73 (2015).

32. Jones, G.E. et al. Microcephaly with or without chorioretinopathy, lymphoedema, or mental retardation (MCLMR): review of phenotype associated with KIF11 mutations. Eur J Hum Genet 22, 881–7 (2014).

33. Tokita, M.J. et al. De Novo Missense Variants in TRAF7 Cause Developmental Delay, Congenital Anomalies, and Dysmorphic Features. Am J Hum Genet 103, 154–162 (2018).

34. Shieh, C. et al. GATAD2B-associated neurodevelopmental disorder (GAND): clinical and molecular insights into a NuRD-related disorder. Genet Med 22, 878–888 (2020).

35. Wang, J. et al. Rett and Rett-like syndrome: Expanding the genetic spectrum to KIF1A and GRIN1 gene. Mol Genet Genomic Med 7, e968 (2019).

36. Chin, C.H. et al. cytoHubba: identifying hub objects and sub-networks from complex interactome. BMC Syst Biol 8 Suppl 4, S11 (2014).

37. Nowakowski, T.J. et al. Spatiotemporal gene expression trajectories reveal developmental hierarchies of the human cortex. Science 358, 1318–1323 (2017).

38. Haney, J.R. et al. Broad transcriptomic dysregulation across the cerebral cortex in ASD. bioRxiv, 2020.12.17.423129 (2020).

39. Willsey, A.J. et al. Coexpression networks implicate human midfetal deep cortical projection neurons in the pathogenesis of autism. Cell 155, 997–1007 (2013).

40. Lukaszewicz, A. et al. G1 phase regulation, area-specific cell cycle control, and cytoarchitectonics in the primate cortex. Neuron 47, 353–64 (2005).

41. Hansen, D.V., Lui, J.H., Parker, P.R. & Kriegstein, A.R. Neurogenic radial glia in the outer subventricular zone of human neocortex. Nature 464, 554–561 (2010).

42. Nowakowski, T.J., Pollen, A.A., Sandoval-Espinosa, C. & Kriegstein, A.R. Transformation of the Radial Glia Scaffold Demarcates Two Stages of Human Cerebral Cortex Development. Neuron 91, 1219–1227 (2016).

43. Lelieveld, S.H. et al. Meta-analysis of 2,104 trios provides support for 10 new genes for intellectual disability. Nat Neurosci 19, 1194–6 (2016).

44. Gillentine, M.A. et al. Rare deleterious mutations of HNRNP genes result in shared neurodevelopmental disorders. Genome Med 13, 63 (2021).

45. Kini, H.K., Silverman, I.M., Ji, X., Gregory, B.D. & Liebhaber, S.A. Cytoplasmic poly(A) binding protein-1 binds to genomically encoded sequences within mammalian mRNAs. RNA 22, 61–74 (2016).

46. Ouellette, J. et al. Vascular contributions to 16p11.2 deletion autism syndrome modeled in mice. Nat Neurosci 23, 1090–1101 (2020).

47. Fiddes, I.T. et al. Human-Specific NOTCH2NL Genes Affect Notch Signaling and Cortical Neurogenesis. Cell 173, 1356–1369 e22 (2018).

48. Consortium, T.S. SPARK: A US Cohort of 50,000 Families to Accelerate Autism Research. Neuron 97, 488–493 (2018).

49. Turner, T.N. et al. Genomic Patterns of De Novo Mutation in Simplex Autism. Cell 171, 710–722 e12 (2017).

50. Manichaikul, A. et al. Robust relationship inference in genome-wide association studies. Bioinformatics 26, 2867–73 (2010).

51. McKenna, A. et al. The Genome Analysis Toolkit: a MapReduce framework for analyzing next-generation DNA sequencing data. Genome Res 20, 1297–303 (2010).

52. Garrison, E. & Marth, G. Haplotype-based variant detection from short-read sequencing. (2012).

53. Rimmer, A. et al. Integrating mapping-, assembly- and haplotype-based approaches for calling variants in clinical sequencing applications. Nat Genet 46, 912–918 (2014).

54. Kim, S. et al. Strelka2: fast and accurate calling of germline and somatic variants. Nat Methods 15, 591–594 (2018).

55. Turner, T.N. et al. Sex-Based Analysis of De Novo Variants in Neurodevelopmental Disorders. Am J Hum Genet 105, 1274–1285 (2019).

56. Xu, X., Wells, A.B., O’Brien, D.R., Nehorai, A. & Dougherty, J.D. Cell type-specific expression analysis to identify putative cellular mechanisms for neurogenetic disorders. J Neurosci 34, 1420–31 (2014).

