## Supplementary material for "Integrated gene analyses of *de novo* mutations from 46,612 trios with autism and developmental disorders": 2.Supplemental_Information

Wang et al.

### Supplemental Information

*Supplemental Information includes Figures S1-S11, Tables S1-S16, and Supplemental References.*

**Figures S1-S11**

Figure S1. Overlap of genes with non-synonymous DNM across phenotype groups.
Figure S2. DNM rate across cohorts.
Figure S3. Significant genes across phenotype by three models.
Figure S4. Significant genes are highly enriched as intolerant genes.
Figure S5. Overlap of NDD-significant genes with reported significant genes.
Figure S6. Brain expression of significant genes.
Figure S7. Expression of NDD and control genes in human cortex.
Figure S8. Expression comparison of identified and reported NDD genes with control genes.
Figure S9. Conditional and unconditional enrichment analyses.
Figure S10. Enriched expression of top genes with sex-biased DNM enrichment significance.
Figure S11. Overlap of significant genes in NDD patients with recalled and no-recall subsets.

**Tables S1-S16** (Note: Tables S3-S8, S10-S16 are provided in spreadsheet)

Table S1. DNM cohorts in this study.
Table S2. DNM rate across cohorts.
Table S3. Integrated coding DNMs.
Table S4. DNM enrichment analysis in siblings (n = 5,241).
Table S5. DNM enrichment analysis in NDD patients (n = 46,612).
Table S6. DNM enrichment analysis in DD patients (n = 31,052).
Table S7. DNM enrichment analysis in ASD patients (n = 15,560).
Table S8. Significance overview of the 615 NDD candidate genes.
Table S9. DNM comparison in ASD and DD patients for the LC615 NDD candidate genes.
Table S10. Intolerance score and DNM in ASD versus DD patients for all significant genes.
Table S11. Genes with significant burden of dnLGD and dnMIS variants in NDD patients (n = 46,612).
Table S12. DNM enrichment analysis in male patients (n = 19,319).
Table S13. DNM enrichment analysis in female patients (n = 8,132).
Table S14. Top hub genes by CytoHubba.
Table S15. DNM enrichment analysis in recalled subset (n = 19,375).
Table S16. DNM enrichment analysis in no-recall subset (n = 27,237).

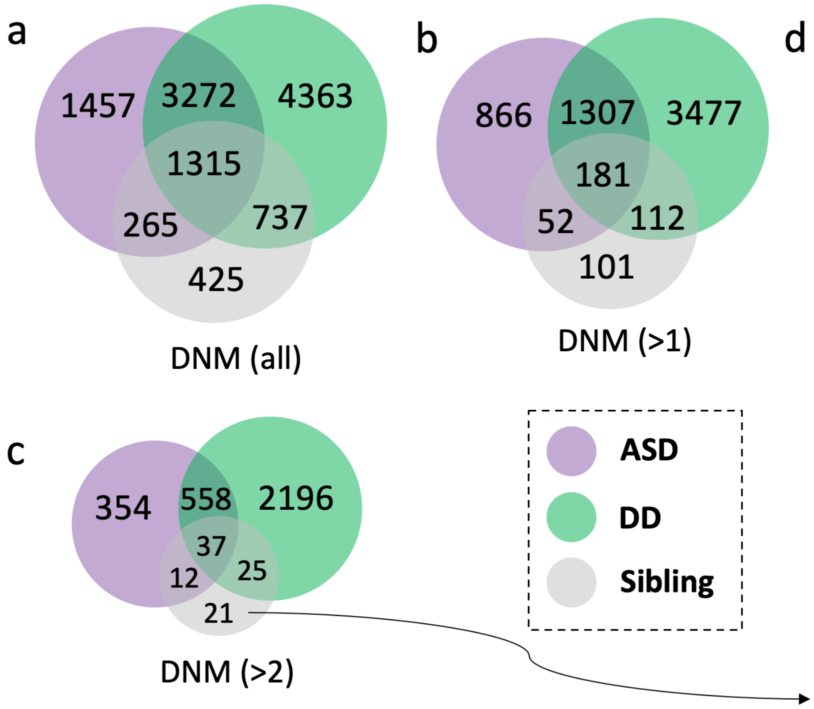


d

DNMs and significance in siblings

| Gene | dnLGD | dnMIS | FDR 5% |
|---|---|---|---|
| *CLEC2D* | 0 | 3 | CH&DR (dnMIS) |
| *CEP20* | 0 | 3 | DR (dnMIS) |
| *FAIM* | 0 | 3 | |
| *ADAM18* | 1 | 2 | |
| *CETP* | 0 | 3 | |
| *CFAP54* | 0 | 3 | |
| *DDX25* | 0 | 3 | |
| *ERBB4* | 0 | 3 | |
| *IPO11* | 0 | 3 | |
| *KLHL21* | 0 | 3 | |
| *MYBL2* | 0 | 3 | |
| *NLRP5* | 0 | 3 | |
| *NMI* | 0 | 3 | |
| *PARD3* | 0 | 3 | |
| *PDZRN3* | 1 | 2 | |
| *RABGAP1* | 1 | 2 | |
| *SPOCK3* | 0 | 3 | |
| *THAP9* | 0 | 3 | |
| *TMCC3* | 0 | 3 | |
| *TRIM17* | 0 | 3 | |
| *YIF1A* | 1 | 2 | |

**Figure S1. Overlap of genes with non-synonymous DNM across phenotype groups.** The majority of genes (82.8%, 2,047/2,472) with DNM in siblings also have DNMs in probands. There are **(a)** 425 genes (all DNMs), **(b)** 101 genes (DNM > 1), and **(c)** 21 genes (DNM >2) with DNMs only in siblings at different filtering level of DNM counts. **(d)** Most of the 21 genes with DNMs (n > 2) exclusively in siblings carry only dnMIS variants, except four genes (*ADAM18*, *PDZRN3*, *RABGAP1*, and *YIF1A*) with one dnLGD variant. Only two genes (*CLEC2D* and *CEP20*) show excess of dnMIS variants in siblings with false discovery rate at 5% (FDR 5%, q-value < 0.05, DNM count > 1).

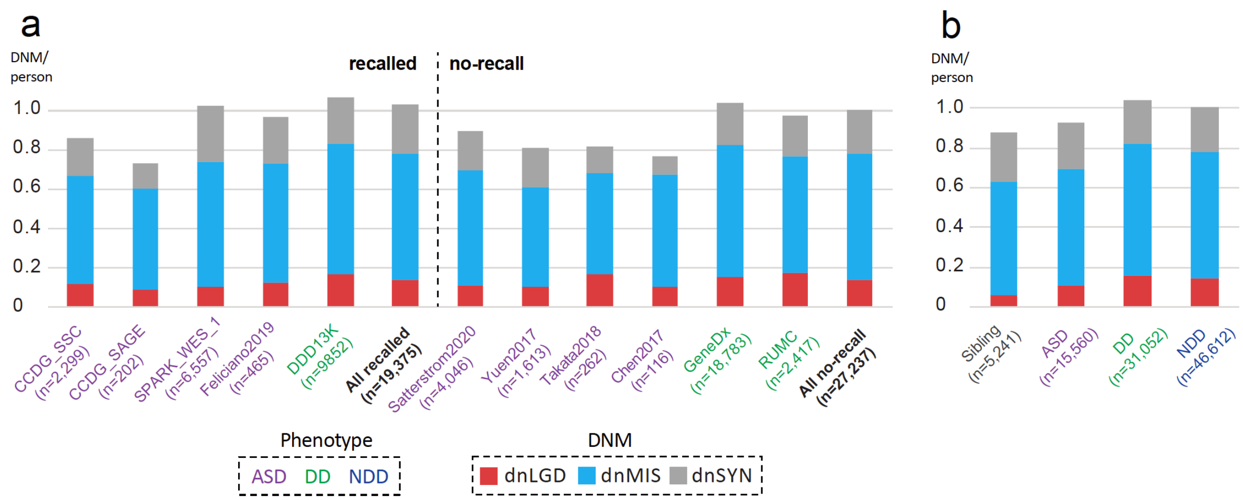


**Figure S2. DNM rate across cohorts.** The number of DNMs per person was calculated for **(a)** each individual cohort and **(b)** each phenotype group after data harmonization with the same filtering criteria applied. The DNM rate is consistent across large cohorts, and also consistent across the cohorts with (recalled) and without (no-recall) reanalysis of the underlying sequencing data. The numbers of probands are also shown with color indicating the phenotype (ASD in purple, DD in green, and NDD in dark blue). Note the bar plot indicates the number of DNMs per person, they are absolute values and no statistics or distributions involved.

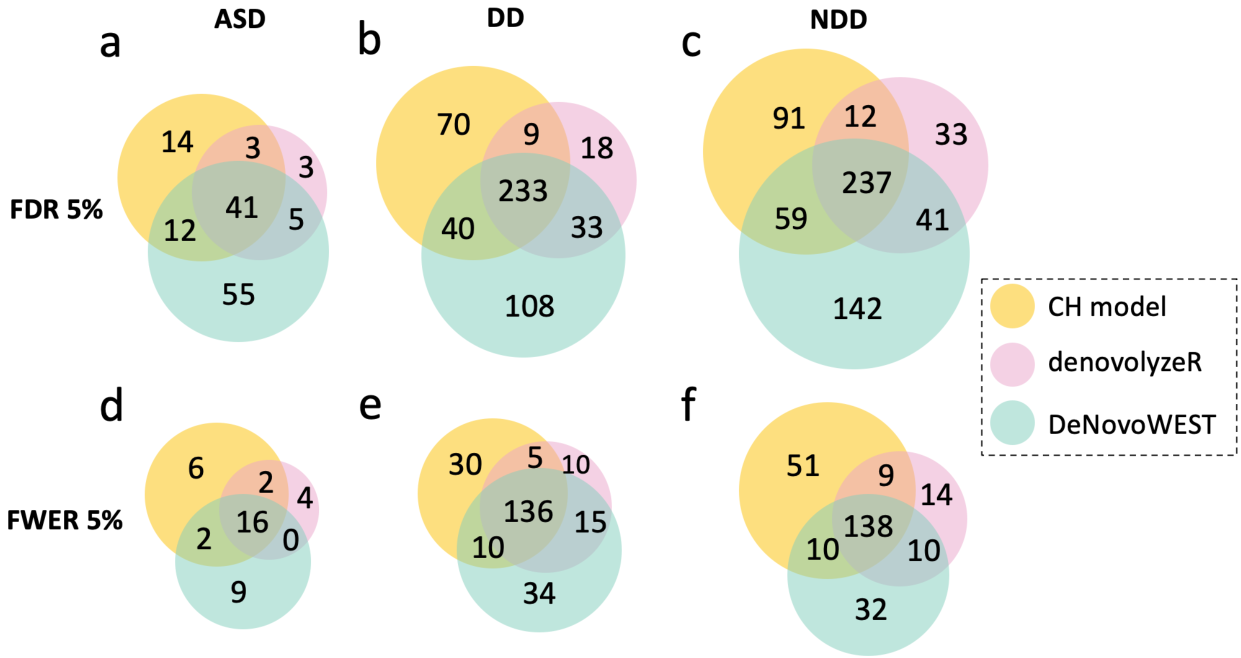


**Figure S3. Significant genes across phenotype by three models.** Overlap of FDR 5% significant genes by three models are shown in **(a)** ASD, **(b)** DD and **(c)** NDD groups in the above panel; similarly, the overlap of FWER 5% significant genes are shown in **(d)** ASD, **(e)** DD and **(f)** NDD groups.

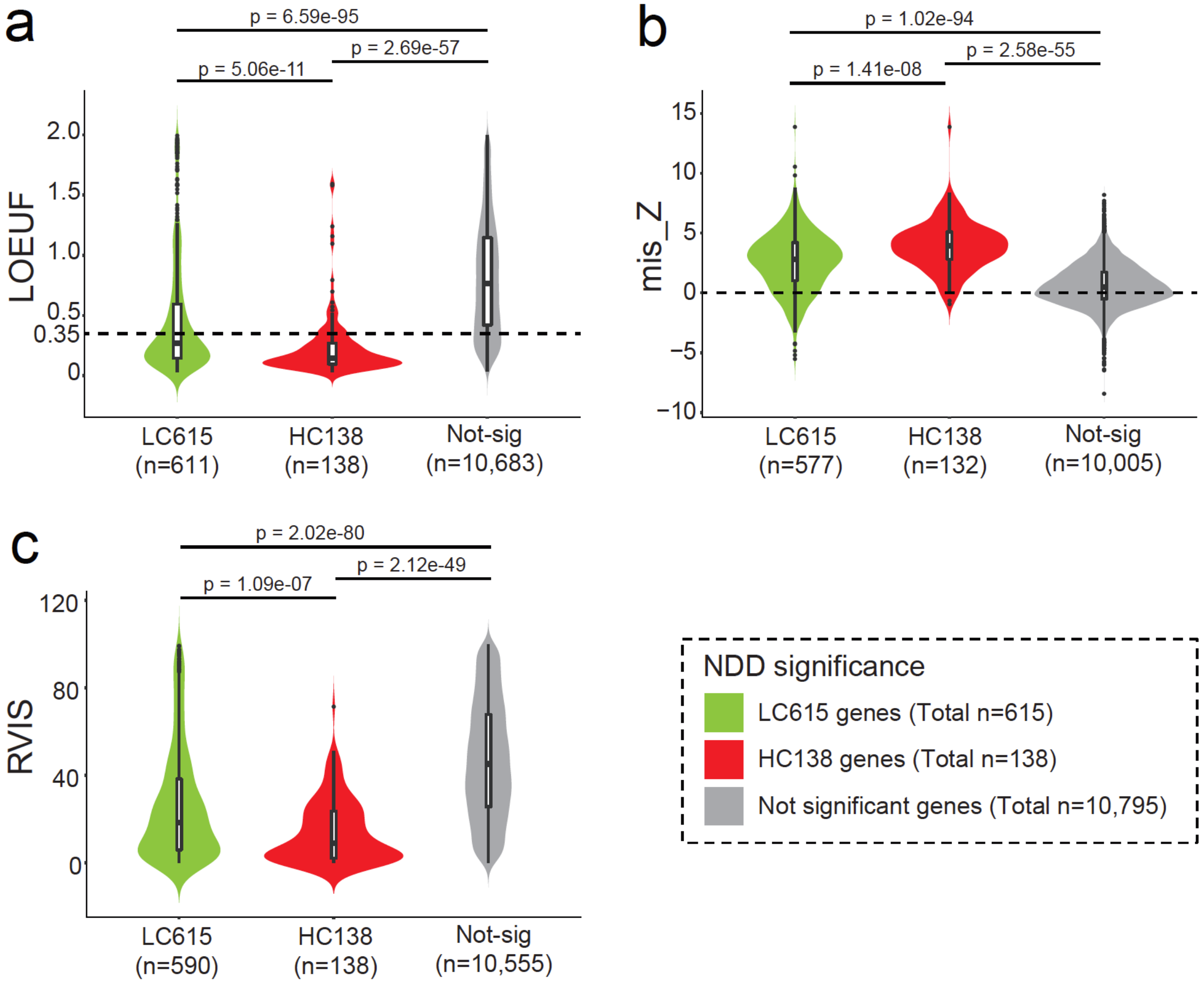


**Figure S4. Significant genes are highly enriched as intolerant genes.** Three intolerant score metrics—**(a)** LOEUF, **(b)** mis_Z, and **(c)** RVIS—were compared across the LC615, HC138, and the rest of gene with DNM in the combined NDD group but not show any DNM significance (Not-sig). The LC615 and HC138 genes identified in this study are significantly intolerant to mutation when compared to the genes not significant, and also between the LC615 and HC138 significant genes, with the corresponding p-value annotated on the top. The number of genes with scores available are shown in parentheses below each category. Wilcoxon two-sample test was performed in R (v3.6.1) with the wilcox.test function. For the box plots, the lower whisker indicates the lowest data point excluding outliers (minima) and the upper whisker indicates the largest data point excluding outliers (maxima); the lower bound indicates the first quartile which is the median of the lower half of the dataset (25th percentile), the upper bound indicates the third quartile which is the median of the upper half of the dataset (75th percentile), and with the middle value of the dataset (50th percentile) indicates in the middle.

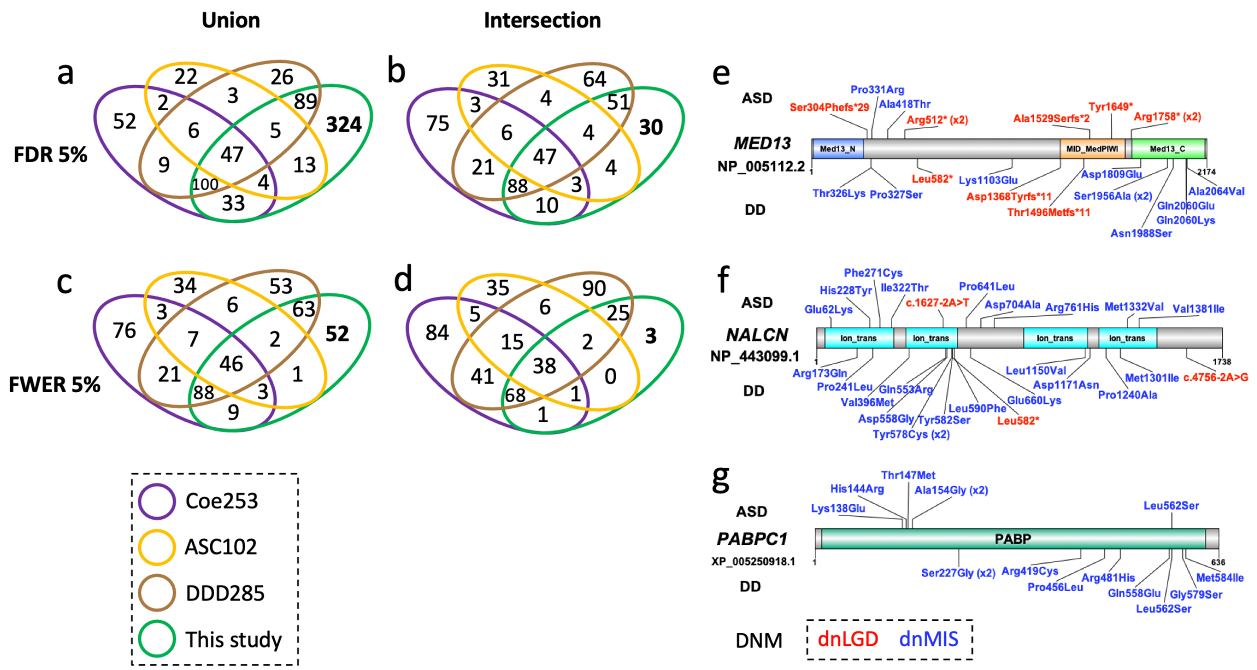


**Figure S5. Overlap of NDD-significant genes with reported significant genes.** Three main published gene lists—Coe253[1], ASC102[2], and DDD285[3]—were considered for the first-step comparison of reported significance. The overlap of these published genes with the genes identified in this study with FDR 5% significance at the **(a)** union set and **(b)** the intersection set of the three models (CH model, denovolyzeR, DeNovoWEST) are shown above; similarly, the overlap with FWER 5% significance in the **(c)** union set and **(d)** intersection set of all three models are shown below. If considered the most stringent of FWER 5% intersection significance, three candidate genes with potential "novel" statistical significance. Protein diagram with dnLGD (red) and dnMIS (blue) variants from ASD (above the diagram, n = 15,560) and DD (below the diagram, n = 31,052) patients for the three candidates are shown in: **(e)** *MED13*, **(f)** *NALCN*, and **(g)** *PABPC1*.

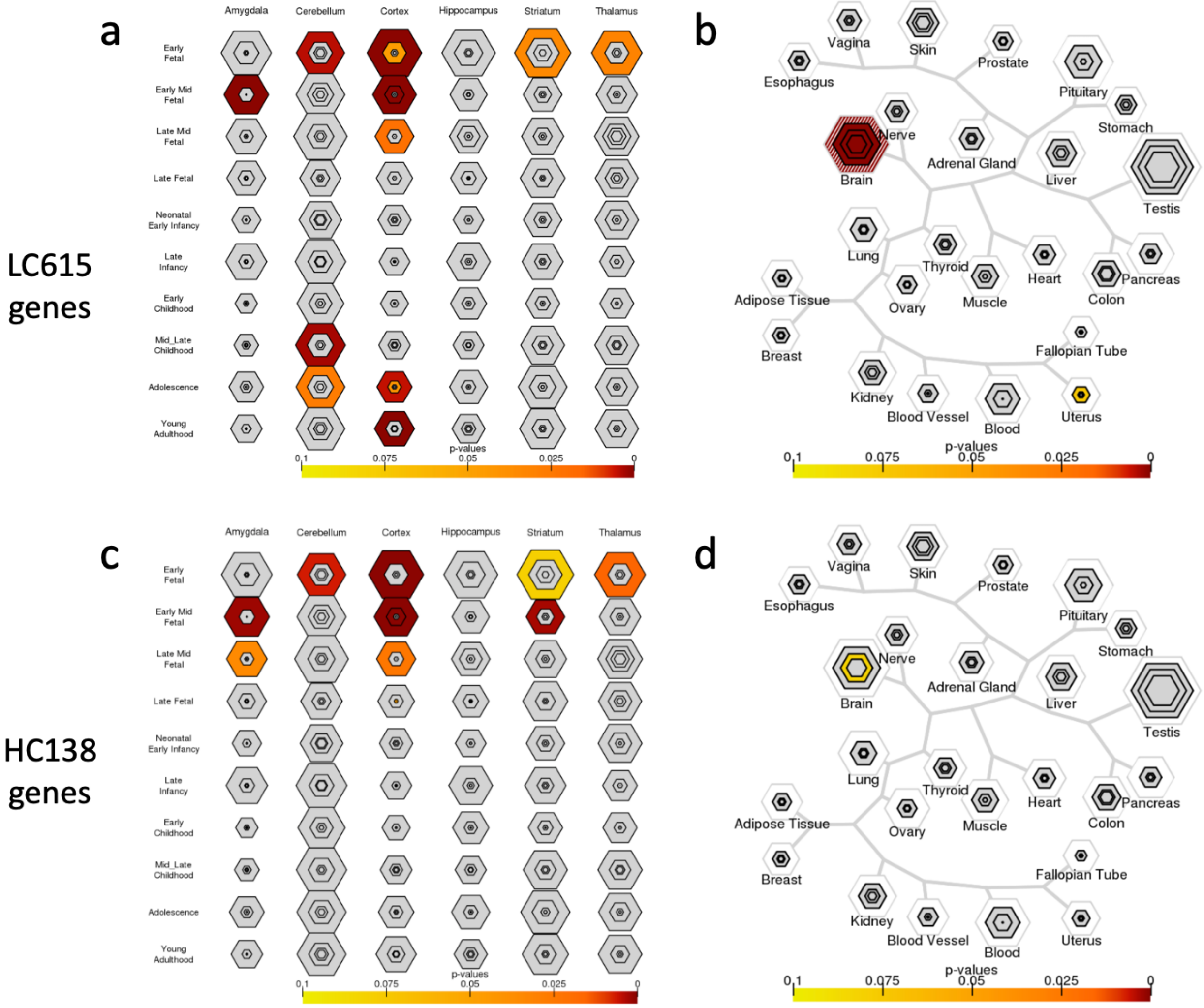


**Figure S6. Brain expression of significant genes. (a)** Specific brain region enrichment during development as determined by CSEA, showing enriched expression of the LC615 genes in the early-mid and late-mid fetal amygdala, early fetal cerebellum, early to late fetal cortex, early to early-mid fetal striatum, and early fetal thalamus. **(b)** Those genes have an enriched expression in the brain as suggested by TSEA. The p-values are from the Fisher's exact test followed by the Benjamini–Hochberg correction. Similar enrichment is also observed for the HC138 genes even with much fewer genes as shown in **(c)** and **(d)**.

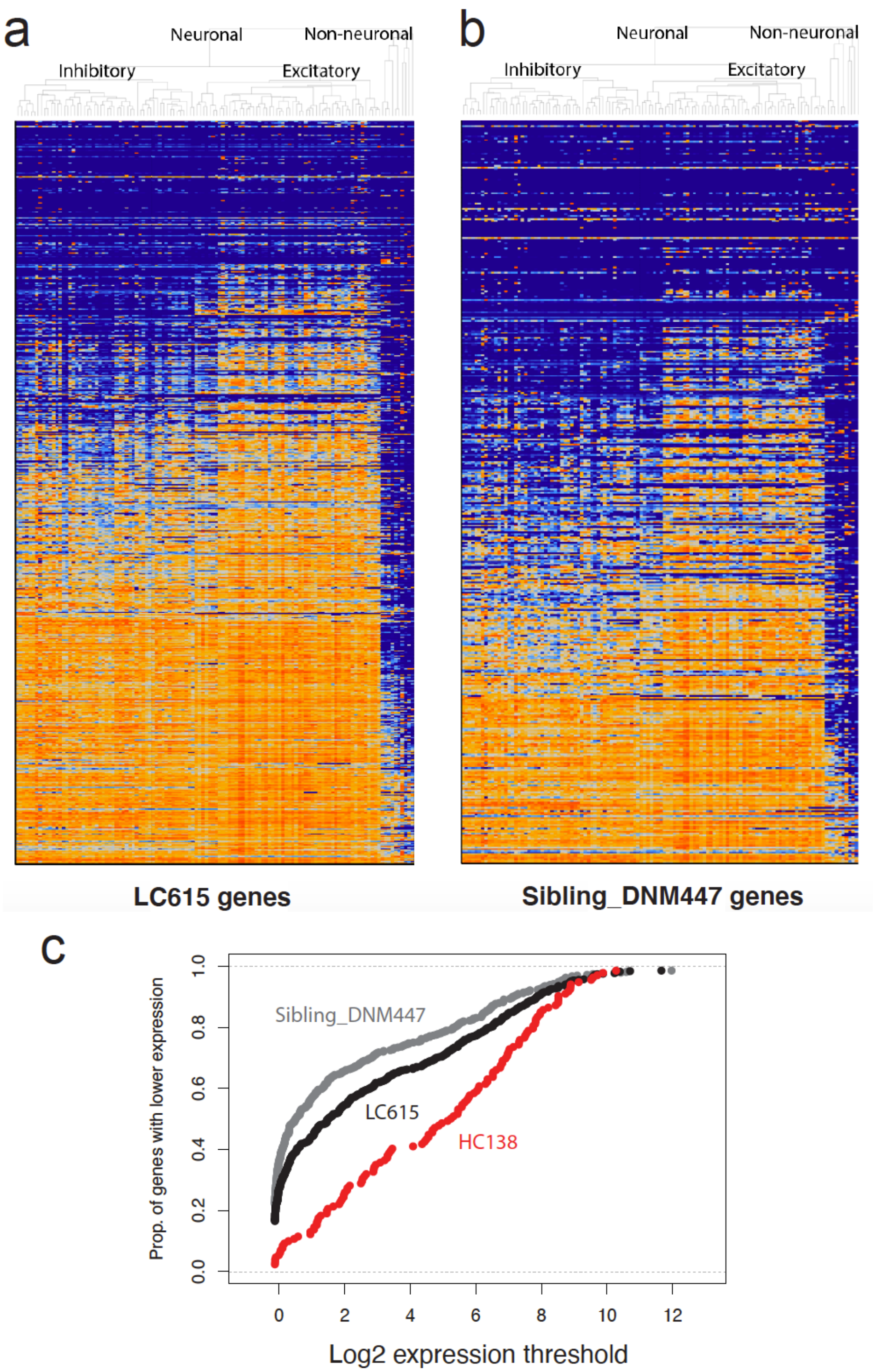


**Figure S7. Expression of NDD and control genes in human cortex.** Expression heatmap of **(a)** LC615 NDD genes and **(b)** 447 sibling DNM (n > 1) genes in 120 cell types identified across six human neocortical areas. **(c)** Empirical cumulative distributions of log2 trimmed mean expression of three gene sets. Rightward-shifted curves indicate more genes with greater expression.

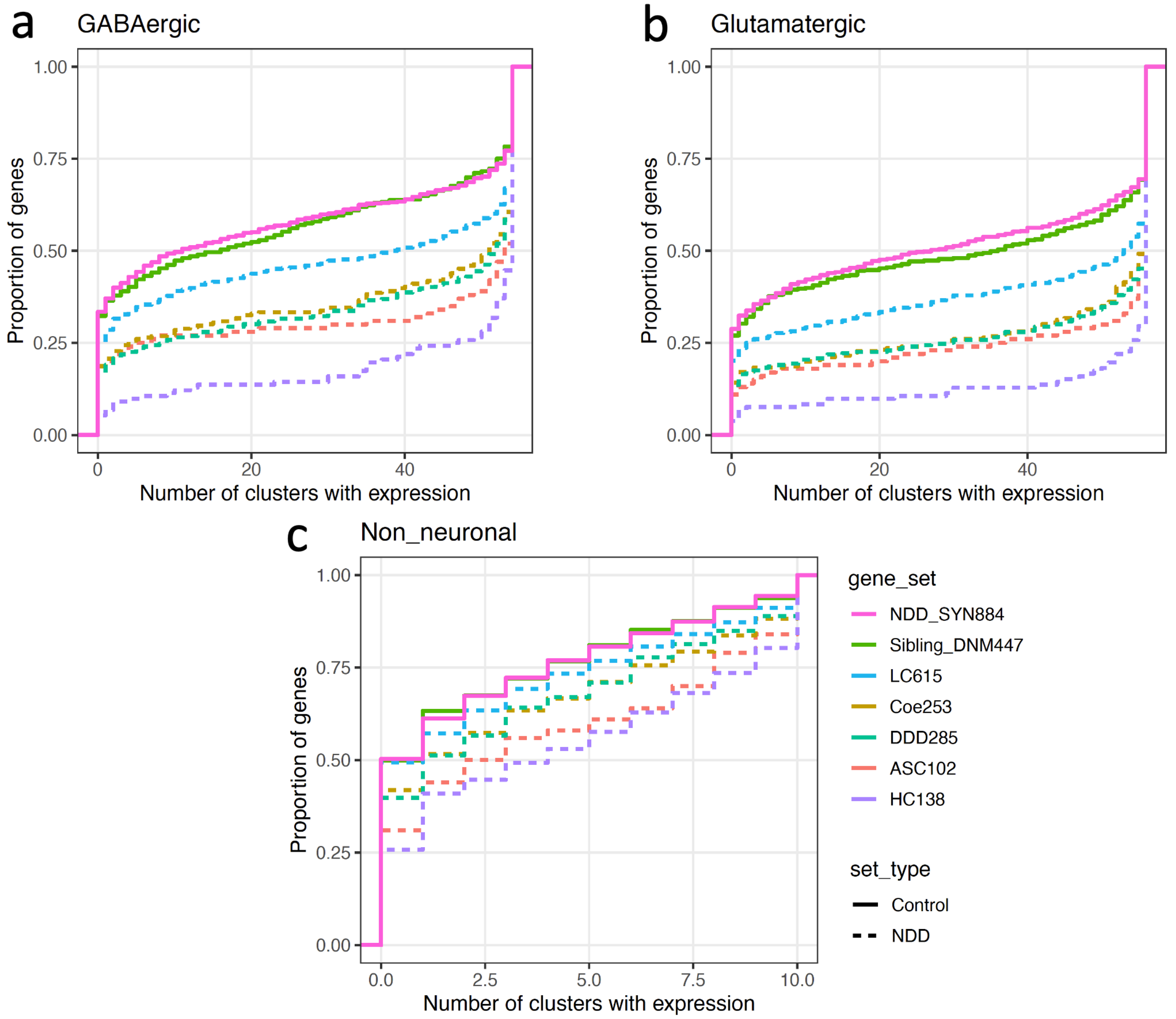


**Figure S8. Expression comparison of identified and reported NDD genes with control genes.** All NDD gene sets are significantly enriched in neuronal types compared to controls. The HC138 genes trend toward more neuronal enrichment than ASC102, although this is not statistically significant. All NDD gene sets (except the LC615 genes) are significantly enriched in non-neuronal cells compared to controls.

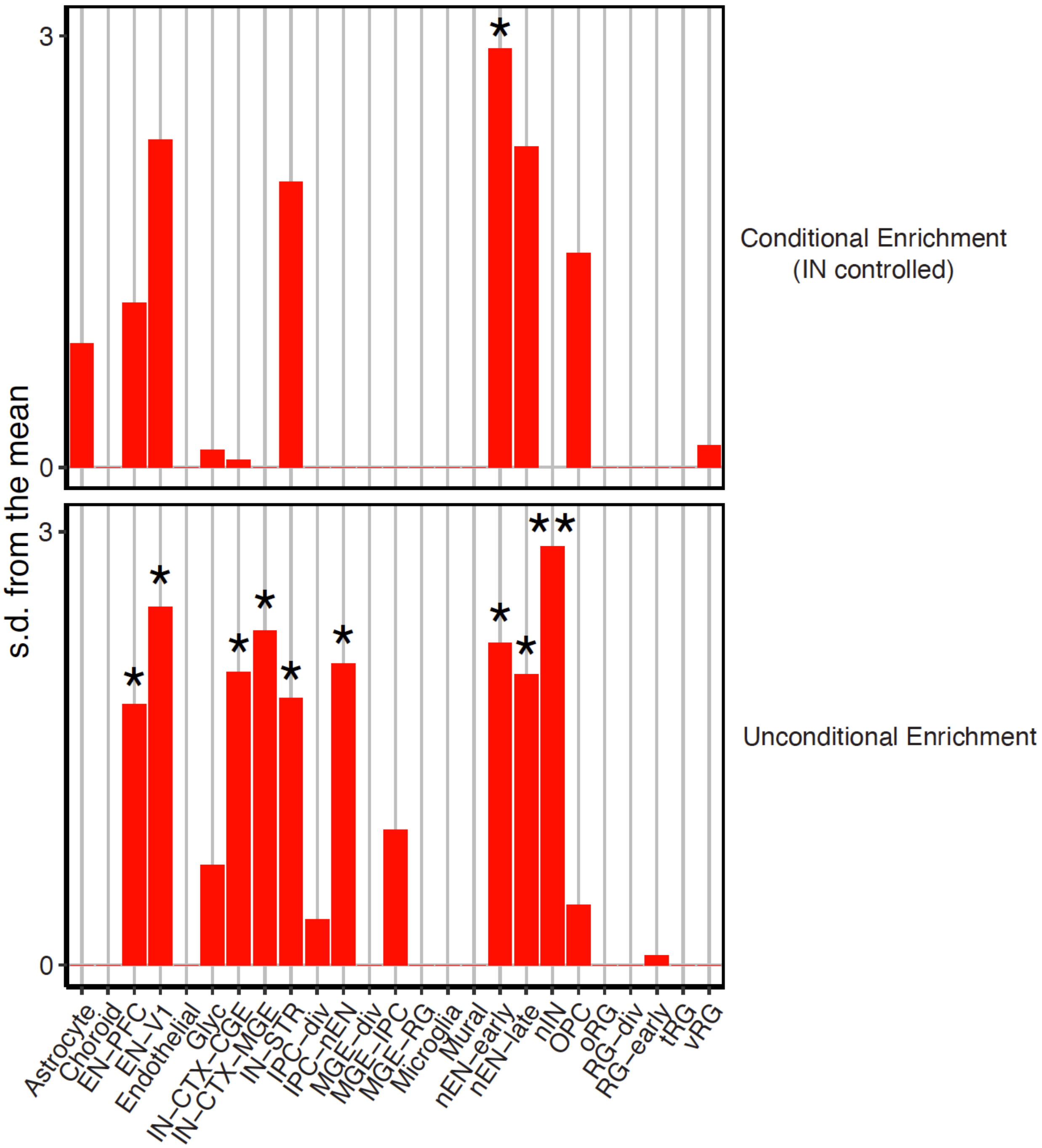


**Figure S9. Conditional and unconditional enrichment analyses.** Conditional enrichment analysis between the neuronal lineage populations found excitatory neurons to have an independent signal with respect to controlling interneuron signal; while neither enrichment nor significance (p = 1 for all) is reached in any neuronal lineage population when controlling for excitatory neurons (* indicates $p < 0.05$, ** indicates FDR $p < 0.05$).

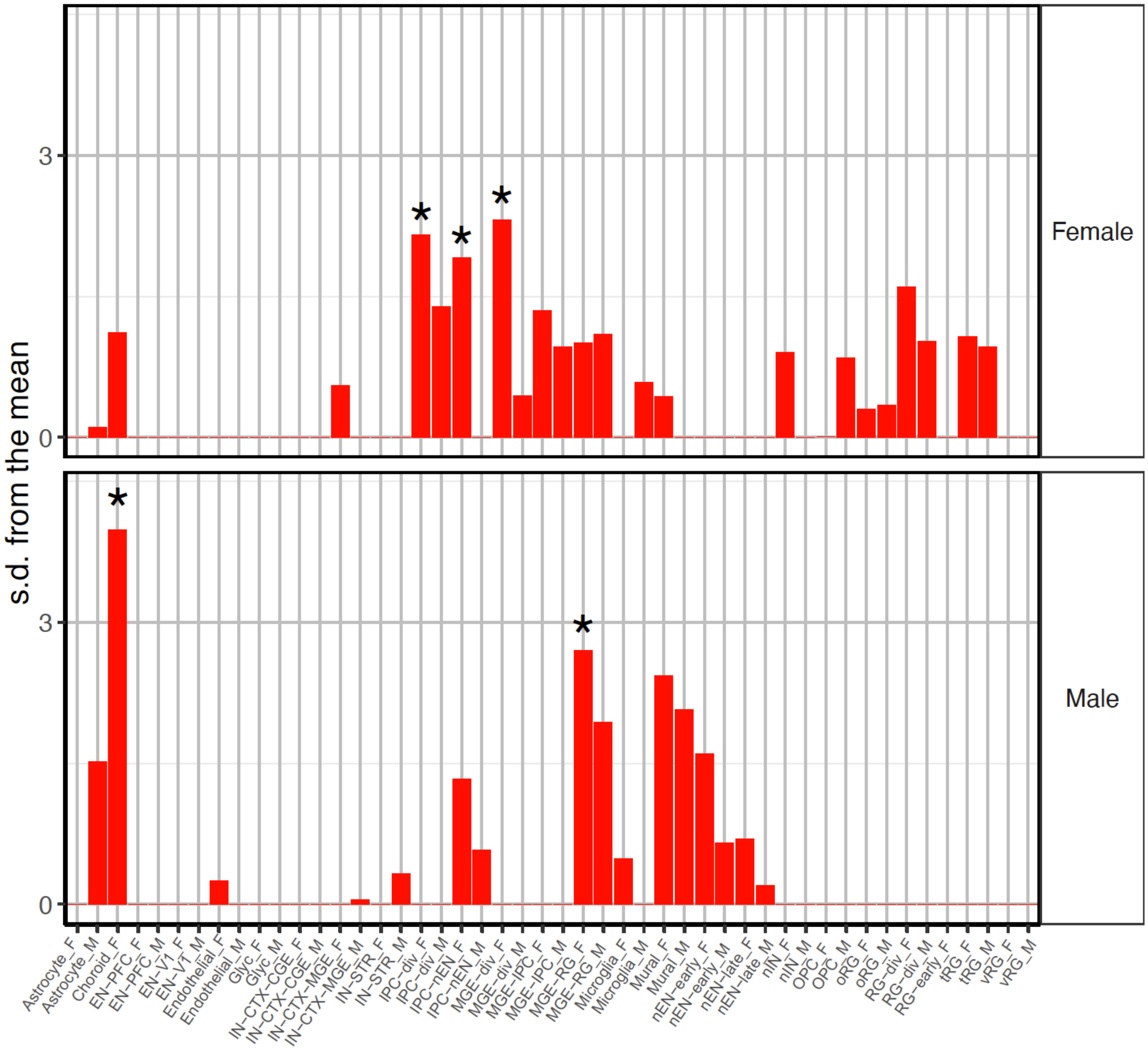


**Figure S10. Enriched expression of top genes with sex-biased DNM enrichment significance.** The top 10 genes for both female and male DNM enrichment were tested with the HC138 genes as background. Enrichment signal was found in the intermediate progenitor cells for the female-enriched genes, and no such signal for male-enriched genes. (* indicates $p < 0.05$).

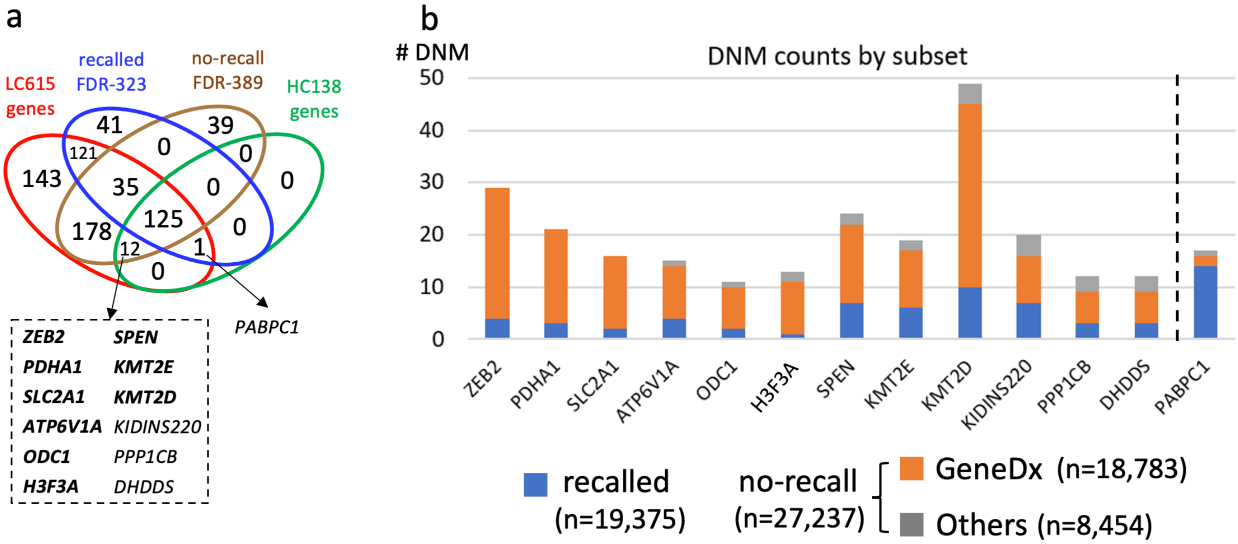


**Figure S11. Overlap of significant genes in NDD patients with recalled and no-recall subsets. a)** Among the HC138 genes in the combined NDD group, there are 12 exclusively significant in the no-recall subset and one gene (*PABPC1*) exclusively significant in the recalled subset. **b)** DNMs were almost exclusively from GeneDx, especially for *ZEB2*, *PDHA1*, and *SLC2A1* where all DNMs are from GeneDx and no DNM from other cohorts in the no-recall subset. Note the bar plot indicates the number of DNMs in each gene, they are absolute values and no statistics or distributions involved.

**Table S1. DNM cohorts in this study.**

| Group | Cohort | Samples | Proband | Male | Female | Sibling | Male | Female | NGS | Individual Sex info | Study |
|---|---|---|---|---|---|---|---|---|---|---|---|
| ASD | *SPARK_WES_1* | *27,256* | *6,557* | *5,229* | *1,328* | *3,034* | *1,551* | *1,483* | *WES* | *Known* | *This study* |
| | SSC | 8,757 | 2,299 | 1,989 | 310 | 1,860 | 879 | 981 | WGS | Known | CCDG_SSC[4] |
| | SAGE | 547 | 202 | 161 | 41 | - | - | - | WGS | Known | CCDG_SAGE[5] |
| | SPARK_pilot | 1,379 | 465 | 377 | 88 | - | - | - | WES | Known | Feliciano2019[6] |
| | MSSNG | 4,174 | 1,613 | 1,268 | 345 | - | - | - | WGS | Known | Yuen2017[7] |
| | ASC | 12,123 | 4,046 | 3,263 | 783 | 347 | 171 | 176 | WES | Known | Satterstrom2020[2] |
| | JASD | 786 | 262 | 191 | 71 | - | - | - | WES | NA | Takata2018[8] |
| | ACE | 348 | 116 | 98 | 18 | - | - | - | WES | NA | Chen2017[9] |
| | **Sub-total** | **55,370** | **15,560** | **12,576** | **2,984** | **5,241** | **2,601** | **2,640** | | | |
| DD | DDD13K | 32,233 | 9,852 | 5,655 | 4,197 | - | - | - | WES | Known | Kaplanis2020[3] |
| | GeneDx | 56,367 | 18,783 | 10,385 | 8,398 | - | - | - | WES | NA | Kaplanis2020[3] |
| | RUMC | 7,254 | 2,417 | 1,377 | 1,040 | - | - | - | WES | Known | Kaplanis2020[3] |
| | **Sub-total** | **95,854** | **31,052** | **17,417** | **13,635** | **-** | **-** | **-** | | | |
| **NDD** | **Total** | **151,224** | **46,612** | **29,993** | **16,619** | **5,241** | **2,601** | **2,640** | | | |

Cohorts are organized into three groups based on primary phenotype (ASD, DD, and NDD). The number of samples in each cohort are total parent–child samples pre data harmonization. The affected (proband) and unaffected (sibling) counts are samples used in the meta-analysis after data QC and harmonization.

**Table S2. DNM rate across cohorts.**

| Subset | Phenotype | Cohort | Proband | dnLGD | rate | dnMIS | rate | dnSYN | rate | DNM | rate |
|---|---|---|---|---|---|---|---|---|---|---|---|
| | ASD | SPARK_WES_1 | 6,557 | 658 | 0.10 | 4,160 | 0.63 | 1,891 | 0.29 | 6,709 | 1.02 |
| | ASD | CCDG_SSC[4] | 2,299 | 261 | 0.11 | 1,273 | 0.55 | 442 | 0.19 | 1,976 | 0.86 |
| **recalled** | ASD | CCDG_SAGE[5] | 202 | 17 | 0.08 | 104 | 0.51 | 27 | 0.13 | 148 | 0.73 |
| | ASD | Feliciano2019[6] | 465 | 58 | 0.12 | 280 | 0.60 | 113 | 0.24 | 451 | 0.97 |
| | DD | DDD13K[3] | 9,852 | 1,624 | 0.16 | 6,571 | 0.67 | 2,296 | 0.23 | 10,491 | 1.06 |
| | | **Sub-total** | **19,375** | **2,618** | **0.14** | **12,388** | **0.64** | **4,769** | **0.25** | **19,775** | **1.02** |
| | ASD | Satterstrom2020[2] | 4,046 | 451 | 0.11 | 2,349 | 0.58 | 814 | 0.20 | 3,614 | 0.89 |
| | ASD | Yuen2017[7] | 1,613 | 160 | 0.10 | 822 | 0.51 | 318 | 0.20 | 1,300 | 0.81 |
| **no-recall** | ASD | Takata2018[8] | 262 | 44 | 0.17 | 133 | 0.51 | 36 | 0.14 | 213 | 0.81 |
| | ASD | Chen2017[9] | 116 | 12 | 0.10 | 66 | 0.57 | 11 | 0.09 | 89 | 0.77 |
| | DD | GeneDx[3] | 18,783 | 2,903 | 0.15 | 12,575 | 0.67 | 3,961 | 0.21 | 19,439 | 1.03 |
| | DD | RUMC[3] | 2,417 | 404 | 0.17 | 1,442 | 0.60 | 507 | 0.21 | 2,353 | 0.97 |
| | | **Sub-total** | **27,237** | **3,974** | **0.14** | **17,387** | **0.64** | **5,647** | **0.21** | **27,008** | **0.99** |
| **All** | **NDD** | **Total** | **46,612** | **6,592** | **0.14** | **29,775** | **0.64** | **10,416** | **0.22** | **46,783** | **1.00** |

The number of DNMs per proband was calculated for each cohort after data harmonization with the same filtering criteria. DNM rate is consistent across cohorts, and also consistent across the recalled and no-recall subset cohorts. This table corresponds to Figure S1.

**Table S9. DNMs comparison in ASD and DD patients for the LC615 NDD candidate genes.**

| Significant genes with non-synonymous DNMs | dnLGD | dnMIS | dnMIS30 | DNMs |
|---|---|---|---|---|
| In ASD only | 42 | 27 | 32 | 13 |
| In DD only | 152 | 158 | 206 | 118 |
| In neither | 245 | 10 | 216 | 0 |
| In both | 176 | 420 | 161 | 484 |
| ASD > DD | 56 | 138 | 58 | 149 |
| ASD < DD | 120 | 282 | 103 | 335 |

Comparison of non-synonymous DNMs, breakdown by mutation class, in ASD (n = 15,560) and DD (n = 31,052) patients was performed for all the LC615 candidate risk genes. For genes with DNM in both ASD and DD patients ("In both"), DNM ratio was further compared with sample size considered for calculation. "ASD > DD" means genes with relatively more DNMs in ASD than DD patients; "ASD < DD" means genes with relatively less DNMs in ASD than DD patients. There are 78.7% (484/615) of genes have DNMs in both ASD and DD samples; more genes have high frequency of DNM in DD (335 genes) than in ASD (149 genes); and more genes have DNMs in DD patients (118 genes) than in ASD patients (13 genes).

### Supplemental References


1. Coe, B.P. *et al.* Neurodevelopmental disease genes implicated by de novo mutation and copy number variation morbidity. *Nature Genetics* **51**, 106-116 (2019).
2. Satterstrom, F.K. *et al.* Large-Scale Exome Sequencing Study Implicates Both Developmental and Functional Changes in the Neurobiology of Autism. *Cell* **180**, 568-584 e23 (2020).
3. Kaplanis, J. *et al.* Evidence for 28 genetic disorders discovered by combining healthcare and research data. *Nature* (2020).
4. Wilfert, A.B. *et al.* Recent ultra-rare inherited mutations identify novel autism candidate risk genes. *bioRxiv*, 2020.02.10.932327 (2020).
5. Guo, H. *et al.* Genome sequencing identifies multiple deleterious variants in autism patients with more severe phenotypes. *Genet Med* **21**, 1611-1620 (2019).
6. Feliciano, P. *et al.* Exome sequencing of 457 autism families recruited online provides evidence for autism risk genes. *NPJ Genom Med* **4**, 19 (2019).
7. RK, C.Y. *et al.* Whole genome sequencing resource identifies 18 new candidate genes for autism spectrum disorder. *Nat Neurosci* **20**, 602-611 (2017).
8. Takata, A. *et al.* Integrative Analyses of De Novo Mutations Provide Deeper Biological Insights into Autism Spectrum Disorder. *Cell Rep* **22**, 734-747 (2018).
9. Chen, R. *et al.* Leveraging blood serotonin as an endophenotype to identify de novo and rare variants involved in autism. *Mol Autism* **8**, 14 (2017).
